# Potential range shifts of the *Anastrepha fraterculus* complex (Diptera:Tephritidae) morphotypes: implications under climate change scenarios

**DOI:** 10.64898/2026.01.08.693998

**Authors:** Damián Freilij, Juan César Vilardi, Paula Gómez-Cendra

## Abstract

The South American fruit fly *Anastrepha fraterculus* is a cryptic species complex that comprises at least eight morphotypes and endangers part of the fruit production across the Americas. In this study, we modeled the current and future potential distributions of the whole complex and each morphotype. Filtered occurrence data (n = 339) were analyzed against six previously selected non-collinear bioclimatic predictors. Ensemble models created with statistical and machine learning algorithms identified ∼6 million km² of environmentally suitable area for the complex, mainly in Brazil and Argentina. Morphotype-level analyses revealed contrasting range extension and suitable areas: Brazilian-1 exhibited the broadest distribution and environmental breadth, whereas Brazilian-2 and –3 were restricted to coastal regions. High-elevation morphotypes (Andean, Ecuadorian, Peruvian, Venezuelan) were strongly constrained by temperature, while the Mexican morphotype distribution was more related with precipitation and displayed a significant potential for expansion into Central and North America. Future projections indicated overall contractions (18–40%) of the areas presenting high-suitability for the presence of the species, along with southward range shifts and increased morphotype overlap in temperate regions; nevertheless, the remaining high-suitable area include the current zones of high production of fruits. These results provide a morphotype-explicit reference for anticipating invasion risk and support area-wide integrated pest management strategies under climate change.

## Introduction

Fruits are a major economic resource for Latin America, accounting for about 25% of the total worldwide production and making the region the largest fruit exporter in the world (FAO, 2019; Garcia, 2024a). Therefore, preventing damage caused by pests is essential for regional economy. One of the most significant threats is represented by fruit flies (Diptera: Tephritidae) that cause substantial economic losses and food waste, often requiring the implementation of costly quarantine restrictions, especially in tropical and subtropical regions (Aluja, 1994; Norrbom, 2010; Araujo *et al*., 2019). Of particular concern are species from the genera *Ceratitis, Anastrepha*, *Rhagoletis*, and *Bactrocera*, which have notable agronomic importance in the region (Garcia, 2024a).

The South American fruit fly, *Anastrepha fraterculus* (Wiedemann), is one of the most polyphagous and widely distributed species in the Americas, infesting over 170 host plants (Hernández-Ortiz *et al*., 2019). Classified as an A1 quarantine pest (EPPO, 2024), it has been reported in the Neotropics, from southern Texas to central Argentina (EPPO, 2024; Hernández-Ortiz *et al*., 2019, Figure 1A). This fly has a patchy distribution following the host availability and suitable weather conditions and it spreads across a remarkable diversity of environments, ranging from sea level to 3000 masl, and from semi-arid regions to high rainfall areas. Its polyphagy, high dispersal capacity and potential local adaptation have enabled this pest to expand its range in a relatively short period (Freilij *et al*., 2024).

**Fig 1.**
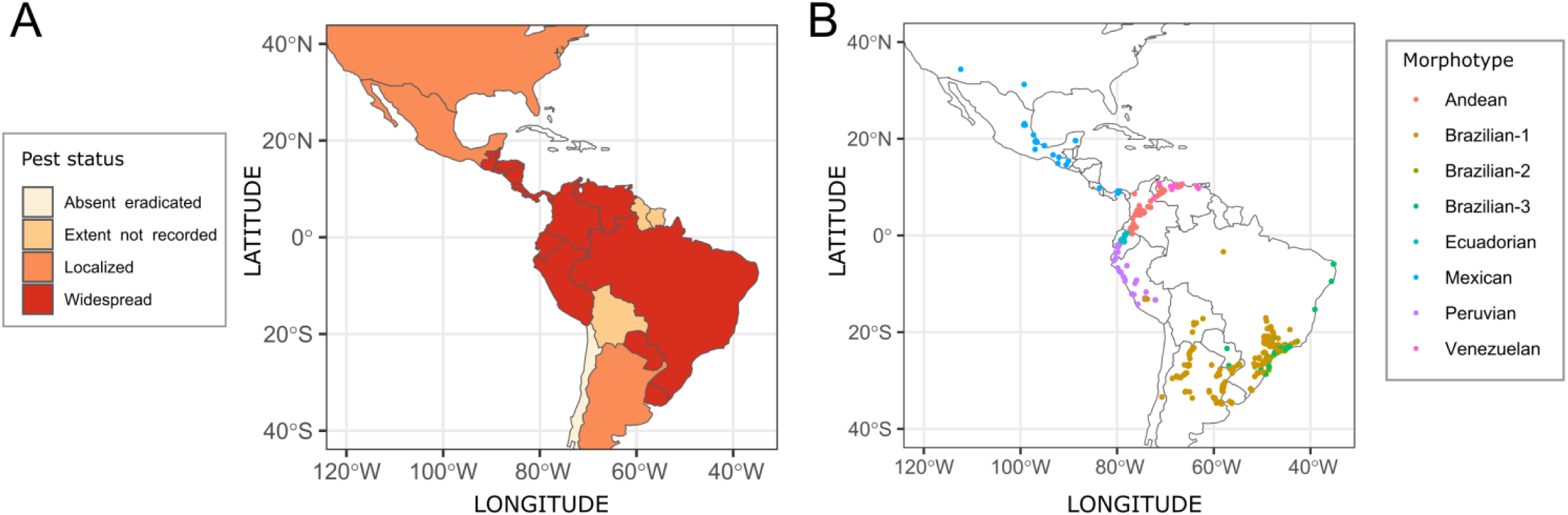
**A**. Pest status report for the *Anastrepha fraterculus* complex per country according to GBIF, EPPO and SpeciesLink databases. **B.** Filtered sampling points (per morphotype) used for species distribution models (see Supplementary Table 1 for location details).

In the past decades, considerable effort has been applied to modeling fruit flies’ current and future distribution by using bioclimatic models. This approach provides each jurisdiction with information to understand the potential invasion hazard and eventually to implement phytosanitary and surveillance measures to mitigate risks (Papadopoulos *et al*., 2024). Given the economic importance of *A. fraterculus* in the region, the application of predictive tools such as species distribution modeling (SDM) to estimate its future distribution is highly relevant. Within this frame a crucial aspect that needs to be addressed is the taxonomic status of *A. fraterculus*. It is currently recognized that this nominal species actually involves an ensemble of cryptic species, known as the *AF* complex (Hernández-Ortiz *et al*., 2012; Selivon *et al*., 2022). The existence of the *AF* complex is supported by multiple lines of evidence, including morphology (Hernández-Ortiz *et al*., 2015, 2019, Prezotto *et al*., 2019), mating compatibility (Vera *et al*., 2006; Rull *et al*., 2013; Dias *et al*., 2016), cytogenetics (Solferini & Morgante 1987; Cáceres *et al*., 2009; Giardini *et al*., 2015; Selivon *et al*., 2022), and molecular analyses (Lopes *et al*., 2013; Sutton *et al*., 2015; Prezotto *et al*., 2019; Freilij *et al*., 2024; Gomulski *et al*., 2024). Morphometric analyses, in particular, have identified eight distinct morphotypes within the *AF* complex: “Mexican” (from Mexico and Central America), “Venezuelan” (lowlands of Venezuela), “Andean” (from the highlands of Colombia and Venezuela), “Peruvian” (lowlands of Ecuador and Peru, *A. sp.4*), “Ecuadorian” (from the highlands of Ecuador and Peru), and three Brazilian morphotypes: “Brazilian-1” (*A. sp.1*, from Brazil and Argentina), “Brazilian-2” (*A. sp.2*), and “Brazilian-3” (*A. sp.3*) (Hernández-Ortiz *et al*., 2015). These morphotypes are adapted to their environments and, for the most part, geographically isolated, with only limited instances of sympatric overlap.

Knowledge of fruit fly ecology and behavior is a key point for effective management (Garcia 2022). Recognizing the cryptic nature of the *AF* complex is essential when designing integrated pest management (IPM) strategies, as differences among morphotypes can influence host preference, dispersal potential, and response to control methods such as parasitoid releases or sterile insect technique (SIT). Incorporating this taxonomic and biological complexity is particularly relevant when applying predictive tools such as SDM, which helps to estimate potential geographic ranges, identify key climate drivers, and anticipate shifts under climate change scenarios. The last few years have seen a surge of studies on SDM and niche models applied to other tephritid species including *Ceratitis capitata* (Szyniszewska *et al*., 2020), *Anastrepha grandis* (Machado Teixeira *et al*., 2022), *A. striata* (Amat *et al*., 2022), *A. suspensa* (da Silva Santana *et al*., 2023), and *A. obliqua* (Santos *et al*., 2019; Batista Degracia *et al*., 2023). They have demonstrated the power of SDMs in revealing potential distributions, understanding climate dynamics, and identifying essential climatic variables with impact on area-wide integrated pest management (AW-IPM). Regarding *A. fraterculus* itself, Santos (2008) addressed ecological zoning for the nominal species in Brazil using a development index for both present and future conditions (for year 2080). These projections predicted a range reduction and that the North, Northwest, and Center-West regions would become entirely unsuitable for the South American fruit fly, while mesothermal conditions would persist in parts of the Southwest and South Brazil despite rising temperatures. These results contrast with a recent work by Gómez-Llano *et al*. (2025), who applied multiple SDM algorithms to assess the potential distributions of the nominal *A. fraterculus* species and two parasitoids that could be used for its biological control. The authors modeled the potential distribution of the species throughout the Americas, under various future climate scenarios and their projections suggested a slight expansion of suitable areas, coupled with a shift toward temperate regions. Moreover, they propose that those changes in suitability may probably lead to an increase in the presence of *A. fraterculus* in the whole region and particularly in Brazil, even though the authors detected some areas of low suitability. Then they stressed that separate analyses for each *A. fraterculus* morphotype are critical to improve prediction accuracy and to adapt IPM strategies to local conditions. On that regard, Selivon *et al*. (2022) had already set an important precedent for the *AF* complex by using SDMs (MaxEnt; Phillips *et al*., 2006) as ecological evidence of cryptic divergence among the three Brazilian morphotypes, identifying key environmental variables and providing useful insights to improve IPM strategies. However, their work focused on the current distribution of the three morphotype and no future projection was modeled.

Building on these previous works, the need arises for further modeling efforts that explicitly incorporate the morphotype variation within *A. fraterculus* and expand it beyond the Brazilian range. The main objective of this study was to model the current and future distribution of the *AF* complex and each of its morphotypes across the Americas. For the current distribution, we aimed to analyze the geographic extent, climatic suitability, and environmental variability associated with each morphotype, as well as to quantify niche similarity and identify potential overlapping areas. Regarding future scenarios, we assessed distributional shifts under different climate projections and identified areas under a significant risk of potential invasion. Overall, we aim to provide an eco-regional and country-level reference, a comprehensive tool to guide and strengthen local and AW-IPM strategies.

## Materials and Methods

### Occurrence data

An extensive search for occurrence data was conducted using the terms “*Anastrepha fraterculus*” and “South American fruit fly” on Google Scholar and PubMed (https://pubmed.ncbi.nlm.nih.gov/) including both research papers and official reports. Additionally, databases such as GBIF (https://doi.org/10.15468/39omei), EPPO (https://gd.eppo.int/), and SpeciesLink (https://specieslink.net) were reviewed to add occurrence points. From these sources, a pest status report map was created. The morphotypes were assigned by direct taxonomic identification when available in the original source or when they belonged to an area where only one morphotype has been observed.

To clean occurrence data, the CoordinateCleaner package (Zizka *et al*., 2019) was used to filter out problematic records based on spatial criteria. Several of the cleaning tests included in the package were applied to remove records that were duplicated, with zero coordinates, outliers or located in capitals or centroids. For reducing spatial bias, occurrence points were subjected to spatial thinning using a minimum distance of 5 km between records, implemented through a manual thinning algorithm. Post-thinning, only data with at least 10 occurrence records were retained for modeling.

### Environmental data

As a first approach, environmental predictors for our study consisted of 19 bioclimatic variables derived from WorldClim 2.1 (Fick & Hijmans, 2017) at a resolution of 2.5 arc minutes. These layers were loaded into the R environment (R Core Team, 2024) and cropped using the stack and crop functions of the raster package (Hijmans, 2015) to a region defined by the extent 30° to 120° West longitude and 40° South 40° North latitude. The coordinates were selected because they encompass the warm regions of the Americas where the *AF* complex can occur. Later, to address multicollinearity and select the most informative predictors, three complementary methods were applied. A principal component analysis (PCA, Hotelling, 1933) was conducted on the scaled variables to identify those with the highest loadings on the first few principal components. Variables that contributed most significantly to the explained variance were retained as candidates for further refinement. In addition, pairwise Pearson correlation coefficients (Pearson, 1896) were calculated using the findCorrelation function from the caret package (Kuhn, 2008), and the environmental variable matrix was pruned to keep only variables with pairwise correlations lower than 0.8. Furthermore, variance inflation factors (VIF, Belsley *et al*., 1980) were computed to identify predictors with redundant information (VIF > 5), and 10 runs of stepwise regressions were employed to refine the selection of variables using the stats package. As for elevation, test runs including this variable did not show significant SDM changes, as it is highly correlated with bioclimatic variables and constant through time, thus it was not included as a predictor for SDM. After SDM, occurrence probability was modeled by elevation for each morphotype and the *AF* complex by means of regressions implemented by geom_smooth (Wickham, 2016).

### Species distribution modeling and niche overlap analysis

To accurately predict a species distribution, it is essential to select the appropriate modeling techniques, as each of them has their own principles and algorithms (Chen *et al*., 2025). Since they offer different advantages and limitations, in this study an ensemble modeling approach was implemented for overcoming the individual models’ limitations and improving the prediction accuracy. The species distribution models (SDMs) were built for both the *AF* cryptic complex, that is, the nominal species *sensu lato*, and for the eight morphotypes separately. With this aim, the biomod2 package was used (Thuiller *et al*., 2009). This modeling platform is widely recognized in this field (Thuiller *et al*., 2009; Bi *et al*., 2013) and its main strength is that it includes around ten different algorithms and allows the user to choose different models, modify their initial conditions, parameters and settings, and integrate them to achieve more robust predictions (Gan *et al*.,2025).

As data only consisted of presence points, pseudo-absence matrices were generated using the BIOMOD_FormatingData function. Multiple strategies were tested, and the random method selected because it provided the most consistent results in preliminary runs. Five replicates of pseudo-absence points were generated using a random strategy, ensuring these points were distributed outside a specified buffer distance from presence locations.

Five alternative algorithms were employed for modeling, encompassing both statistical and machine learning approaches: Generalized Linear Models (GLM), Generalized Boosted Models (GBM), Random Forests (RF), Maximum Entropy (MaxEnt), and Generalized Additive Models (GAM). For GAM, the GAM_mgcv algorithm was used with s_smoother for flexible smoothing, while MaxEnt (Phillips *et al*., 2006) employed 200 iterations per run with 10.000 background points, using automatic selection of linear, quadratic, product, threshold, and hinge features, a regularization multiplier set to the default (1), and a prevalence of 0.5 to optimize species-environment relationships. Models were calibrated by cross-validation with a training set of 70% of the data (considered a conservative option) while the remaining 30% was used for testing the reliability of the models. Five replicate runs were conducted to account for variability arising from random data partitioning, utilizing the bigboss optimization strategy. Additionally, variable importance was assessed across five iterations.

Model performance was assessed using the True Skill Statistic (TSS), ROC AUC (Receiver Operating Characteristic Area Under the Curve) and Cohen’s Kappa (KAPPA, Cohen, 1960). The evaluation results were summarized using the bm_PlotEvalBoxplot function, and to further understand the influence of environmental predictors on morphotypes distribution, response curves were generated using the bm_PlotResponseCurves function. Ensemble models were generated by combining predictions from individual models above a 0.7 metric selection threshold using three different aggregation methods: weighted mean (based on performance metrics), arithmetic mean, and median. The consistency among these approaches was checked to confirm coherence in the final outputs.

Distribution models were developed under current climatic conditions (climate data for 1970-2000) and projected for two future scenarios with contrasting trajectories to assess potential shifts in species distributions under diverse climate futures, spanning the periods 2021–2040 and 2041–2060. Projections were based on two global climate models: Meteorological Research Institute – Earth System Model version 2, MRI-ESM2-0 (Yukimoto *et al*., 2019) under the Shared Socioeconomic Pathways (SSP; IPCC 2021) SSP3-7.0 scenario, representing a high radiative forcing pathway with moderate-to-high socioeconomic challenges (moderate), and UK Earth System Model, UKESM1-0-LL (Williams *et al*., 2018) under the SSP5-8.5 scenario, which assumes high greenhouse gas emissions and minimal mitigation efforts (pessimistic). The previously selected bioclimatic variables were downloaded from WorldClim for each scenario and processed as stated above.

Additionally, Schoener’s D (Schoener, 1968) and the *I* statistic (Warren *et al*., 2008) were used to quantify niche overlap between species. These indices measure the degree of similarity between ecological niches, assessing how much the distributions of two taxonomic units align in space. Both metrics were computed for all pairwise morphotype combinations using the raster.overlap function of the ENMTools package (Warren *et al*., 2021), and results were visualized in a heatmap generated by ggplot2.

### Range shifts and alpha diversity maps

Binary ensemble maps (presence-absence) were generated using a cutoff value of 0.7, which corresponded to the threshold maximizing the True Skill Statistic (TSS) for the ensemble model. These maps were used to analyze range shifts for each morphotype and the whole *AF* complex under current and future climatic scenarios. Thus, areas classified as presence correspond to regions with suitability values above 0.7, representing zones of high predicted occurrence where the most drastic distributional changes are expected. Raster layers for current and future scenarios were imported and processed using the terra package (Hijmans, 2021). To estimate the potential range shifts, the BIOMOD_RangeSize function was employed. A stacked map of range changes was created to visualize spatial patterns using rasterVis::levelplot. For computing the surface area of raster cells across space, by conversion of pixels to km^2^, cellSize and global functions of the terra package were employed.

Alpha diversity was calculated and mapped by summing the binary raster layers representing morphotypes presence/absence in a given area for each scenario. This approach quantifies the morphotype richness (the number of morphotypes present) at each grid cell, providing a direct measure of the species local biodiversity and its changes across time.

## Results

### Occurrence data

A total of 805 geographical records were retrieved throughout the Americas for the *AF* complex, and only one of them could not be assigned to a morphotype (Supplementary Table 1 for detailed information on occurrence points). *A*fter filtering and spatial thinning, a total of 339 points remained for the *AF* complex (Table 1, Supplementary Figure 1). They spread along 15 countries, and their elevations range from 0 to 4500 masl (Supplementary Figure 1). The number of valid records for modeling morphotypes ranged from 11 to 165 (Figure 1B, Table 1), reflecting disparities in occurrence data and/or the amount of research studies for each one. Unfortunately, the Venezuelan and Ecuadorian morphotypes records were particularly scarce, thus caution is advised when interpreting their SDM results.

**Table 1:**
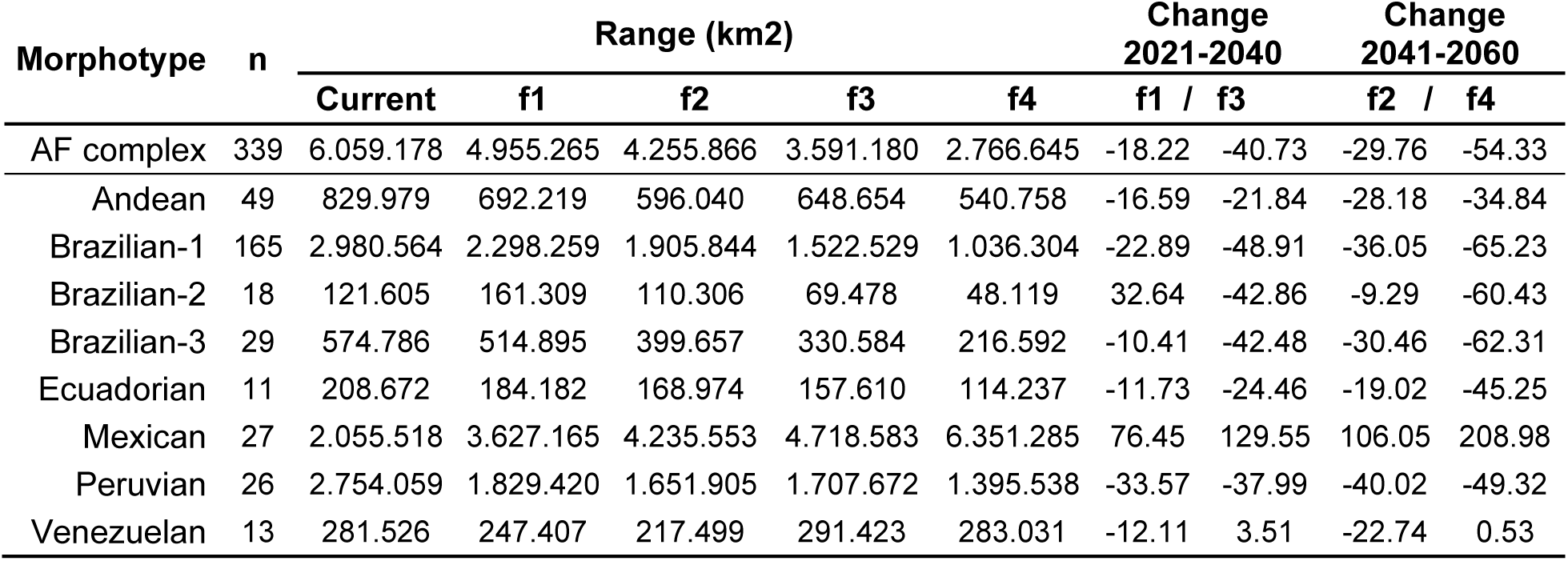
Potential distribution areas (km^2^) per morphotype and for the whole *AF* complex across different time periods and scenarios. Areas correspond to regions with high suitability values (>0.7 based on binary ensemble maps). n= number of records; f1= MRI-ESM2-SSP3-7.0 for 2021-2040; f2= MRI-ESM2-SSP3-7.0 for 2041-2060, moderate scenario; f3= UKESM1-0-LL-SSP5-8.5 for 2021-2040; UKESM1-0-LL-SSP5-8.5 for 2041-2060, pessimistic scenario. The change is expressed as a percentage and represents the range between the MRI-ESM2-SSP3-7.0 and UKESM1-0-LL-SSP5-8.5 scenarios relative to the current distribution.

### Environmental data

The multistep selection process resulted in six bioclimatic variables with low multicollinearity and high explanatory power after modeling: BIO1 (Annual Mean Temperature), BIO2 (Mean Diurnal Range), BIO8 (Mean Temperature of Wettest Quarter), BIO10 (Mean Temperature of Warmest Quarter), BIO13 (Precipitation of Wettest Month), and BIO14 (Precipitation of Driest Month) (Table 2, Supplementary Figure 2).

**Table 2.** Mean percentage of bioclimatic variable importance in the ensemble potential distribution model for each taxonomic unit. BIO1 = Annual Mean Temperature, BIO2 = Mean Diurnal Range, BIO8 = Mean Temperature of Wettest Quarter, BIO10 = Mean Temperature of Warmest Quarter, BIO13 = Precipitation of Wettest Month, BIO14 = Precipitation of Driest Month.

| **Morphotype** | **BIO1** | **BIO2** | **BIO8** | **BIO10** | **BIO13** | **BIO14** |
| --- | --- | --- | --- | --- | --- | --- |
| *AF* complex | 45.5 | 22.2 | 19.7 | 8.25 | 2.2 | 2.1 |
| Andean | 25.4 | 12.6 | 18.3 | 28.8 | 11.4 | 3.7 |
| Brazilian-1 | 39.0 | 15.5 | 9.9 | 24.7 | 2.9 | 7.9 |
| Brazilian-2 | 23.1 | 21.4 | 17.6 | 22.8 | 10.7 | 4.3 |
| Brazilian-3 | 20.5 | 42.3 | 11.7 | 8.14 | 10.1 | 7.3 |
| Ecuadorian | 14.1 | 8.6 | 6.1 | 67.3 | 1.6 | 6.1 |
| Mexican | 16.2 | 9.6 | 8.2 | 14.7 | 32.0 | 19.3 |
| Peruvian | 23.5 | 11.8 | 12.8 | 34.4 | 11.2 | 6.3 |
| Venezuelan | 29.8 | 2.9 | 1.7 | 23.8 | 25.3 | 16.5 |

### Species distribution modeling and niche overlap analysis

Pseudo-Absences points, model evaluation metrics and response curves per algorithm and morphotype or complex are shown in Supplementary Figures 3-11, as well as evaluation metrics, response curves, and the contribution of each algorithm to the ensemble model. As a general pattern, Random Forest (RF) consistently performed as the most robust and reliable algorithm across all metrics (KAPPA, ROC AUC, and TSS), with high medians (0.96, 0.95, and 0.92, respectively) and minimal variability (±0.2 SD). Generalized Linear Model (GLM) also showed a good performance, yielding strong results across the board, though with slightly higher variability (±0.3 SD) compared to RF. MAXENT demonstrated moderate performance, with lower medians than RF and GLM (KAPPA=0.89; ROC= 0.95; TSS=0.84), along with greater variability. In contrast, GAM and GBM exhibited the poorest calibration results overall, characterized by significantly lower medians (KAPPA≈0.9) and higher dispersion (±0.5 SD) (Supplementary Figures 3-11). Differences between weighted mean, median and arithmetic mean were minimal (data not shown), and the arithmetic mean projections are presented.

The ensemble model for the *AF* complex was in agreement with the current distribution reports and showed an extensive potential distribution area throughout the tropics and subtropics (Fig. 2A). The extent of the climatically suitable area for *A. fraterculus* under current climate conditions (calculated from the binary 0.7 TSS threshold projection, *i.e*., suitability ≥ 70 %, representing high suitability) was ∼6.000.000 km^2^ (Table 1). The countries with the highest percentage of highly suitable areas were Brazil (39.72%), Argentina (18.88%), USA (6.94%), Peru (6.38%) and Mexico (3.95%) (see Supplementary Table 2 for percentages details per country and ecoregion). From an environmental perspective, the cryptic species complex encompasses several ecoregions with contrasting characteristics (Cerrado, Dry Chaco, Humid Pampas, moist forests; Supplementary Table 2), and exhibits a bimodal occurrence probability for elevation, with a peak around 1000 masl and another at 3000 masl (Fig. 2B). The most important variables for the *AF* complex distribution were Annual Mean Temperature, Mean Diurnal Range, and Mean Temperature of Wettest Quarter (Table 2, Supplementary Figure 3).

**Fig 2.**
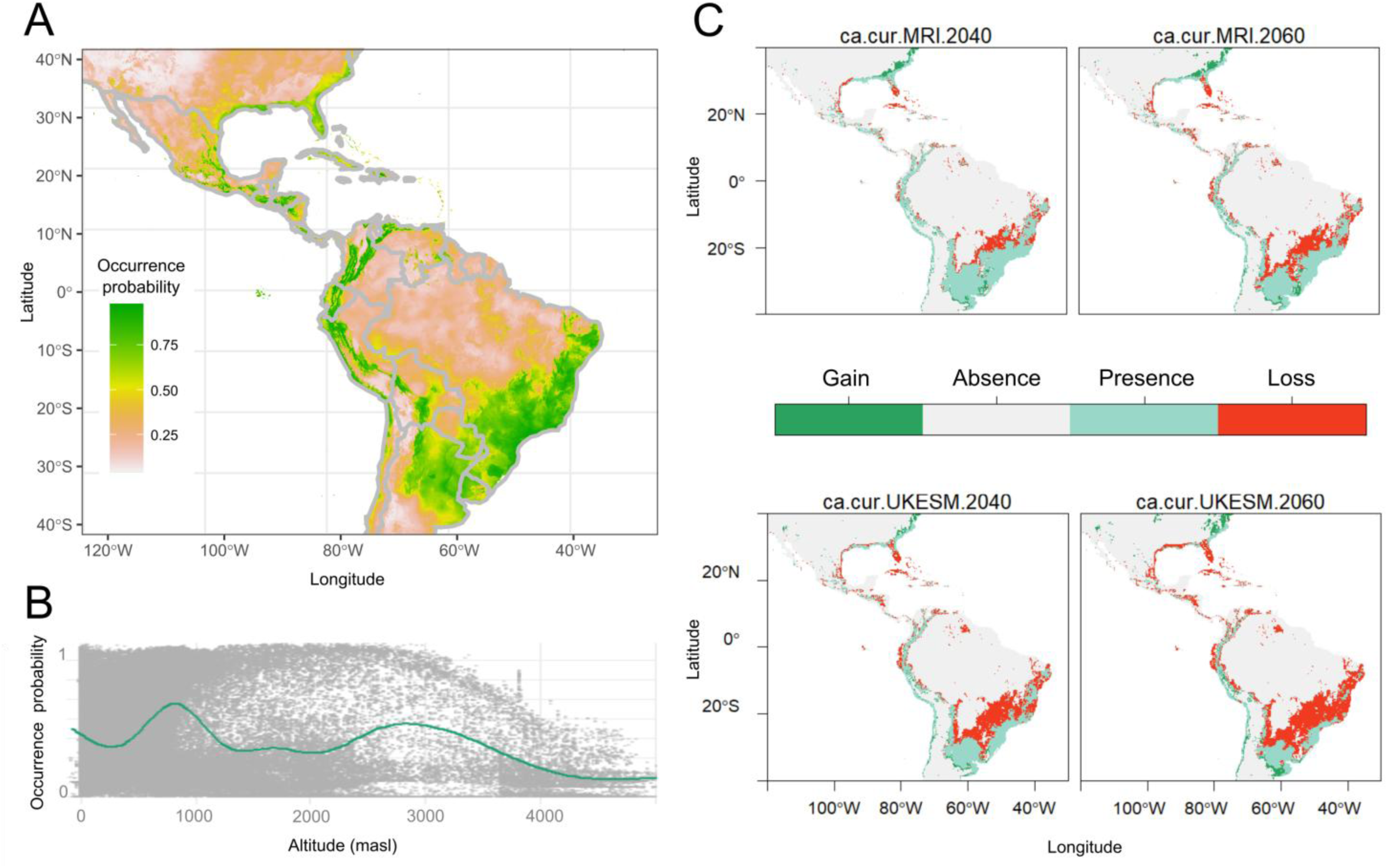
**A**. Current potential distribution of the *AF* complex in the Neotropical Realm. **B.** Probability of occurrence as a function of elevation by a GAM regression. **C.** Range change areas (ca) under different possible future climate scenarios: MRI-ESM2-SSP3-7.0 for 2021-2040 and 2041-2060 (MRI, moderate scenario, top panels); UKESM1-0-LL-SSP5-8.5 for 2021-2040 and 2041-2060 (UKESM, pessimistic scenario, bottom panels). Areas correspond to regions with suitability values above 0.7, based on binary ensemble maps.

Regarding the morphotypes current potential distributions, our results highlighted specific geographical patterns, as well as contrasting ecological niches (Figure 3A; see Supplementary Table 2 for country and ecoregions percentages, and Supplementary Figures 3-11 for modeling details). The Brazilian-1 morphotype exhibited the highest extension and wider climatic niche among morphotypes (Figure 3A), with a potential area of almost 3.000.000 km^2^ (high suitability ≥ 0.7) of which 47.34% corresponded to Brazil, and 23.16% to Argentina (Supplementary Table 2). More than 40% of its total area comprised ecoregions as the Cerrado, Alto Paraná, Atlantic forests, Uruguayan savanna and Araucaria moist forests (Supplementary Table 2), and the morphotype exhibited the same bimodal occurrence probability with elevation as the *AF* complex, one peak around 1000 masl and other at 3000 masl (Supplementary Figure 12). As for Brazilian-2 and Brazilian-3, ranges were significantly narrower (∼120.000 and 575.000 km^2^ respectively) and mainly restricted to Brazilian coastal areas (Figure 3A) with Annual Mean Temperature and Mean Diurnal Range as the predictors with the highest importance (Table 2). The occurrence probability for the Andean, Ecuadorian, Peruvian and Venezuelan morphotypes was higher in highland zones, around 3000 masl, between 12°N – 15°S latitude, 70°W – 80°W longitude and they are distributed in the Northern Andes and Central Andes uppermost portion (Figure 3A, Supplementary Figure 12). The Andean morphotype presence was higher in the Occidental, Central and Oriental valleys of Colombia, and extended northwards up to the Lake Maracaibo region, and southwards to central highlands of Ecuador. The most relevant variables for its distribution were Annual Mean Temperature and Mean Temperature of Warmest Quarter (Table 2, Supplementary Figure 4). The Ecuadorian morphotype exhibited a similar but narrower (∼200.000 km^2^) potential distribution, with a stronger effect of Mean Temperature of Warmest Quarter. Regarding the Peruvian morphotype, most of its distribution encompassed the northwest portion of the coastal, highland and rainforest regions of Peru, and the coastal and sierra regions of Ecuador. The Venezuelan morphotype presented a narrow distribution, ∼280.000 km^2^ along northern Venezuela, the Oriental cordillera valleys of Colombia, central Ecuador and northwestern Peru, mainly influenced by precipitation and Annual Mean Temperature variables (Table 2). Regarding the Mexican morphotype, 20.37% of its potential area corresponded to Mexico and 2.55% to the USA, where the morphotype is actually present nowadays (Figure 3A, Supplementary Table 2), but results pointed out that 33.70% of its current plausible distribution was located in the Cerrado region of Brazil, which showed high potential establishment risk due to niche similarity. The most important variables for its distribution were Precipitation of the wettest and driest month (Table 2).

**Fig 3.**
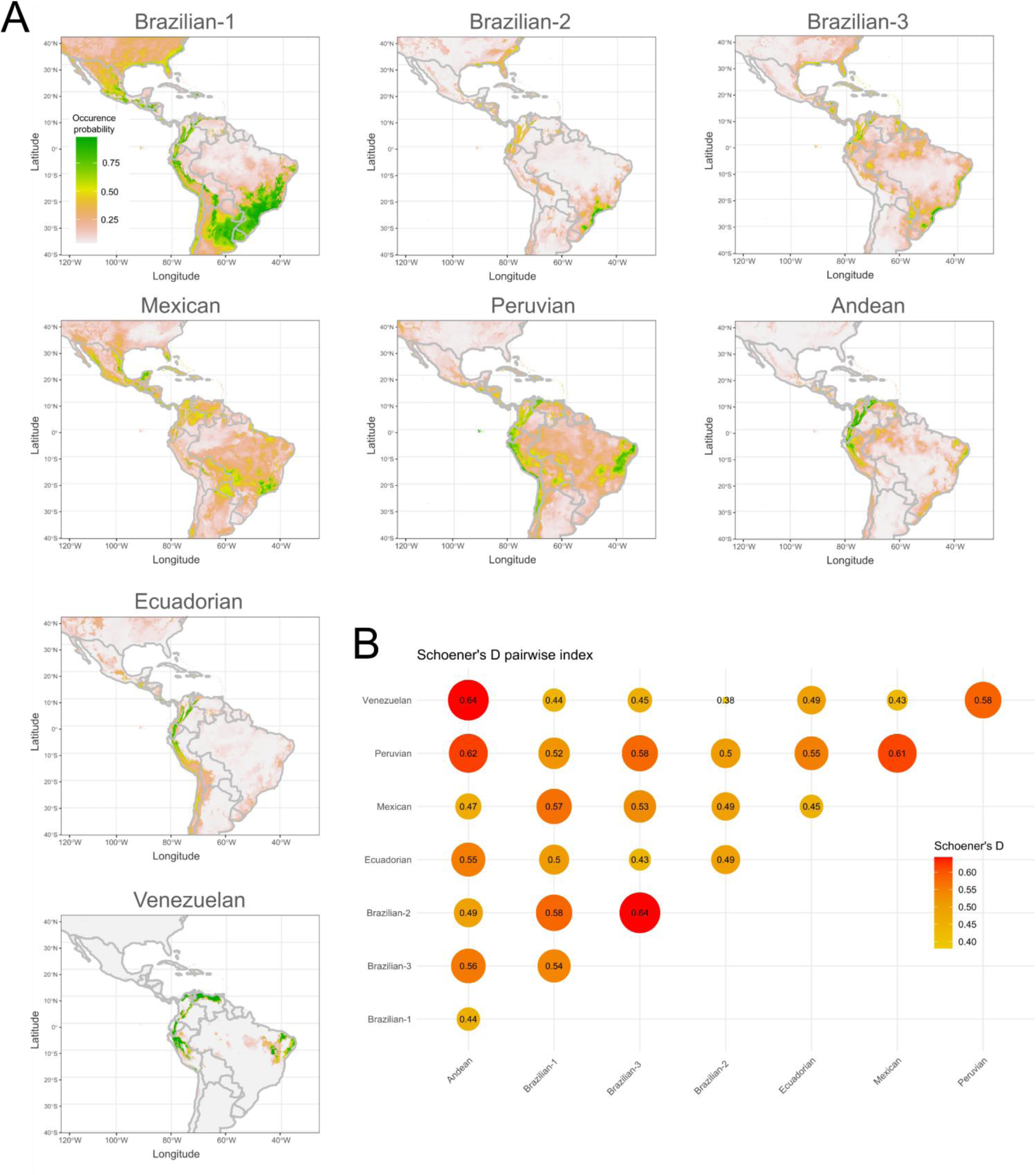
**A**. Current potential distribution of the *AF* complex morphotypes. **B.** Shoener’s D pairwise heatmap for those morphotypes.

Complementing the individual results of the SDM, Schoener’s D pairwise analysis revealed the highest degree of niche identity for the pairs Andean-Venezuelan, Brazilian-1 and –2 and Peruvian-Andean (Figure 3B). Additionally, the analysis indicated significant niche similarity between Peruvian and Brazilian-3, Venezuelan and Peruvian, as well as Mexican-Peruvian and Mexican with Brazilian-1, despite these groups not currently being sympatric. The lowest niche identity was found between the pairs: Venezuelan-Brazilian-2; Ecuadorian-Brazilian-3; Andean-Brazilian-1; Ecuadorian-Mexican (Figure 3B). *I* statistic results yielded the same pattern, but with higher values (data not shown).

### Range shifts and alpha diversity maps

The comparison among current and future projections revealed a negative range shift for the *AF* complex, with an overall reduction of regions with high suitability (≥ 0.7) between ∼18 and 40%, equivalent to 1.103.913 and 2.467.998 km^2^ of its potential area for the period 2021-2040 (Figure 2C, Table 1). Unsurprisingly, the strongest effect was under UKESM1-0-LL-SSP5-8.5, a pessimistic scenario. Despite the reduction in areas, this contraction would not disrupt the connectivity among the major fruit-producing regions of South America. The country with the highest *AF* complex range reduction for 2021-2040 was Brazil (∼30 to 60%, Supplementary Table 2) followed by Mexico (∼30 to 45%). On the other hand, future distribution projections pointed to a potential displacement of high suitability areas to southern latitudes, with an increase in the Pampas ecoregion of Argentina, southwest Uruguay, northern and central Chile. An increase was also observed for the South Atlantic region of the USA (Figure 2C, Supplementary Table 2). About Bolivia a significant reduction between 26-35% for 2021-2040 and 35-60% for 2041-2060 was predicted, with a migration to the south. For Ecuador the coastal area would experience an occurrence reduction, and for Peru a more modest decline and shift in the northern and southern regions were detected (Figure 2C, Supplementary Table 2).

Regarding the different *AF* complex morphotypes, future projections revealed notable changes in high climatic suitability zones (≥ 0.7 range change maps per morphotype are shown in Supplementary Figures 3-11). The Andean morphotype was projected to experience a total reduction of approximately 16-21% between 2021 and 2040, and 28-35% between 2040 and 2061 (Table 2). Despite these reductions, connectivity among the identified areas in Colombia, Venezuela, and Ecuador is expected to be maintained and even increased (Supplementary Figure 4). The Ecuadorian morphotype is expected to undergo a similar reduction, with a decline of approximately 11% and 24% during the same periods, while maintaining connectivity among its current areas and not significant changes in Ecuador (Supplementary Figure 5, Supplementary Table 2). Regarding the Peruvian morphotype, even with an overall reduction of 33.57 to 37.99%, under the UKESM1-0-LL-SSP5-8.5 scenario a small increase for Argentina and Chile was predicted (Supplementary Table 2). The Venezuelan morphotype did not show a clear change along time (Table 2, Supplementary Figure 7). The prediction for Brazilian-1 is an overall decrease in all scenarios –22.89 to –48.91% for 2021-2040 and –36.05 to – 65.23% for 2041-2060 (Table 2, Supplementary Figure 8, Supplementary Table 2), specially for inland Brazilian Atlantic dry forests, and an increase for the Humid Chaco and Northeast Brazil restingas regions. As for Brazilian-2 the predictions were inconclusive in Brazil (Supplementary Figure 9, Supplementary Table 2), as the overall trend for different scenarios was not clear (Table 2). On the other hand, the Brazilian-3 morphotype would decrease its current distribution in Northern Brazil (Supplementary Figure 10, Supplementary Table 2), while the suitable areas in Argentina and Uruguay could significantly increase. The Mexican morphotype range was expected to expand its high suitability areas in all scenarios, increasing its area by ∼15-55% in Mexico, with a displacement of 86080 –158591 km2 to the South Atlantic USA region. In Guatemala and Nicaragua its suitable area was predicted to increase 17.93-48.93% and 60-137% of its current distribution respectively.

Spatial morphotype richness (i.e. morphotype overlapping) is depicted in alpha diversity maps (Figure 4) and it revealed that currently, one-quarter of the potential colonized area is shared, with half of this overlap involving only one other morphotype. The two main regions of potential sympatry –Southeast Brazil and the Northern Andes-are projected to persist across all future scenarios, while the likelihood of sympatry in regions like Northern Brazil is expected to decline (Figure 4).

**Fig. 4.**
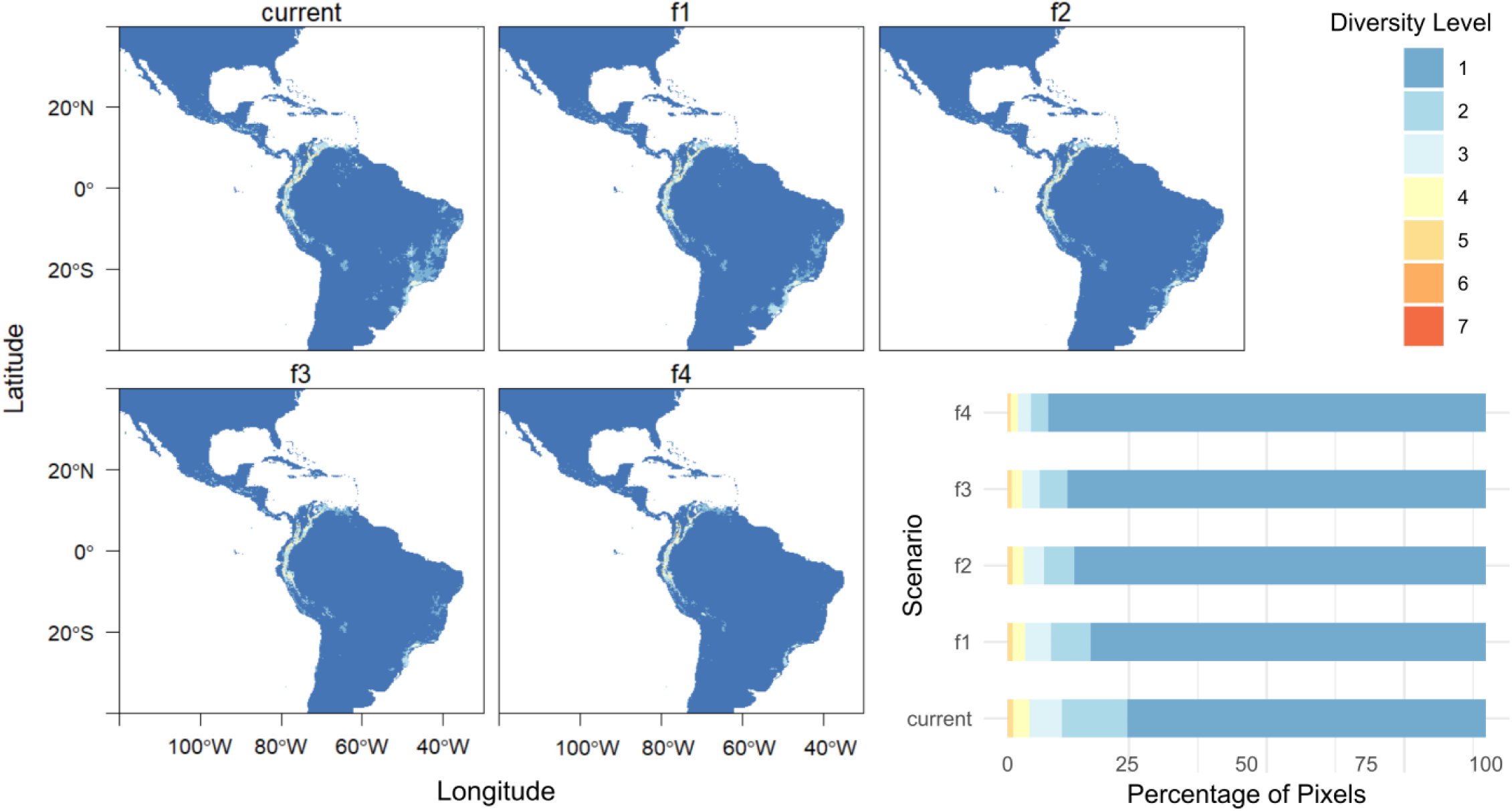
Alpha diversity maps for current and possible future climate scenarios. f1= MRI-ESM2-SSP3-7.0 for 2021-2040; f2= MRI-ESM2-SSP3-7.0 for 2041-2060; f3= UKESM1-0-LL-SSP5-8.5 for 2021-2040; UKESM1-0-LL-SSP5-8.5 for 2041-2060. Diversity values are based on binary ensemble maps with suitability > 0.7. Barplots represent the percentage of diversity levels > 0 per scenario.

## Discussion

Comprehensive knowledge of pest species biology is crucial for designing effective integrated pest management (IPM) strategies and promoting agroecosystems restoration. The focus of our study was the South American fruit fly *Anastrepha fraterculus,* a major fruit pest.

Broadly speaking, fruit flies are more affected by temperature, humidity, and light (Garcia, 2009), but the finer details can be widely variable. For example, Vera *et al*. (2002) found that the distribution of *Ceratitis capitata* in the Andes was limited by low maximum temperatures more than by the absolute minimum temperature, as was commonly believed. Meanwhile, in hot areas like California, the suitability for the same species was dependent on the humidity and therefore irrigation programmes had a significant effect (Szyniszewska *et al*., 2020). Precipitation and humidity can have drastically different effects on different species or areas. For example, the chance of droughts made precipitation on the warmest quarter the most relevant variable for *A. grandis* (Machado Teixeira *et al*., 2022) but rainfall did not affect the adults of *A. ludens* and *A. obliqua* in semi-field conditions (Montoya *et al*., 2009). However, the following generation can be affected, as proved by several studies where precipitation was positively or negatively correlated with the population size (Abad *et al*., 2018) or the number of generations per year (Machado Teixeira *et al*., 2022). It is clear that researching local environmental patterns is essential because distribution and genetic variability of flies populations are strongly associated with temperature, precipitation and soil moisture. While laboratory studies help identify physiological constraints for a species, its actual distribution in the wild would reflect the combined effect of macro– and micro-environmental factors in addition to seasonal fluctuations (Vera *et al*., 2002). Such knowledge is crucial to understand population dynamics and improve the capacity of predicting the response to climate change.

*A. fraterculus*, like others of the same genus, is considered an opportunistic and highly polyphagous species with the potential to adapt to different environments, all traits that explain how it became one of the major pests affecting fruit production in the Americas. Additionally, it represents a complex of cryptic species (*AF* complex) with genetic, behavioral and ecological divergence (Selivon *et al*., 2022). It has been proposed (Freilij *et al*., 2024) that the rapid cryptic divergence of the *AF* complex would have involved ecological niche diversification resulting in local adaptation. Therefore, studying the effect of both regional and local environmental patterns can provide significant insight into the crucial factors that drive the ecology and dynamics of the pest at different scales (Aluja *et al*., 2012). Species Distribution Modeling (SDM) is a relatively new and innovative tool both for ecology and IPM. SDM allows the prediction of species geographic distribution based on occurrence and different environmental variables data, helping to define the potential niche for a species. In turn, this enhances the understanding of the ecological requirements of species and the impacts of climate change, providing valuable information for conservation strategies and pest control (Guisan & Thuiller, 2005).

SDM has grown considerably in recent years, driven by software that simplifies model building (Kass *et al*., 2024). Yet, key challenges remain: selecting appropriate variables, acknowledging model assumptions, balancing algorithmic complexity, and ensuring data quality and model robustness. Packages like biomod2 address these issues (Thuiller, 2014), and therefore it was selected for its capacity to overcome individual algorithm limitations through more robust ensemble models. In this study, five algorithms were implemented, combining 339 occurrence points with six environmental variables. The ensemble model was rather robust, especially for Receiver Operating Characteristic Area Under the Curve (ROC AUC), with values larger than 0.8 even for the metric that showed the poorer results (Kappa) for most of the morphotypes and surpassing 0.6 in all cases. Potential current distribution was modeled, and predictions were projected for two scenarios and time periods. Previous SDM efforts had been restricted either to the nominal species in Brazil (Santos, 2008), to the three Brazilian morphotypes within that country (Selivon *et al*., 2022), or more recently, to the nominal species at a continental scale (Gómez-Llano *et al*., 2025), but without considering morphotypes variability. Despite these valuable contributions, a large-scale projection including all morphotypes and extending into future scenarios was still lacking.

Our study, as the first to model the current and future potential distribution of all morphotypes, contributes to understanding the ecological factors shaping the dynamics of the complex. Consequently, it can also help to predict the response of the different morphotypes to climate changes and could be used as a guide to select areas where surveillance and controls should be increased. Therefore, the results can be used as reference material by governments and similar institutions that need to take informed decisions. Nevertheless, a caveat is in order. Significant asymmetries persist among morphotypes: information is particularly scarce for the Venezuelan and Ecuadorian morphotypes and for *AF* populations in Bolivia and inaccessible areas from Brazil. These gaps reflect broader disparities in scientific capacity and historical sampling biases, underscoring the need to expand systematic records and strengthen collaboration. From a management standpoint, such limitations hinder effective monitoring and prevention, as poorly documented populations may go unnoticed until reaching advanced stages of establishment or spread. A noteworthy initiative addressing this challenge is the recent cooperation between more than 60 researchers in compiling the volume *Management of Fruit Flies in the Americas* (Garcia, 2024b) which covers both general aspects of fruit fly management and detailed IPM strategies for 18 countries in the Americas. When combined with SDM inferences of ecological drivers, suitable area estimations, and physiological experiments, such initiatives can significantly strengthen the assessment and implementation of strategies for fruit fly management in the Neotropics.

As a contribution to this goal, the present study showed *AF* broad potential distribution area throughout the tropics and subtropics, spanning most of South America, Central America and the southernmost portion of North America. This extensive area of ∼6.000.000 km^2^ across contrasting characteristics suits the opportunistic and high adaptability nature of the species, which diverged into different morphotypes in a relatively short period of time (Freilij *et al*., 2024). The areas deemed as highly suitable are in agreement with those previous studies assessing geographical range by other methodologies. Santos (2008) employed ecological zoning in Brazilian territories and found medium suitability for inland zones, and higher for the Southeast (eastern São Paulo, southern Minas Gerais), Northeast (coast of Bahia), Center-West (south-central Mato Grosso do Sul), and North (Acre, Rondônia, southwestern Amazonas, northwestern Pará) of Brazil. Alongside this, the approach taken by Selivon *et al*. (2022) considered the three Brazilian morphotypes and found that *A. sp.*1 was widely distributed and extended to higher elevations further from the equator and the coast, as well as to drier locations. In contrast*, A. sp*.2 and *A. sp*.3 were more similar in geographic and bioclimatic variables, with more limited distributions. Despite using different bioclimatic predictors and suitability criteria, our results showed similar current distribution patterns. Gómez-Llano *et al*. (2025) recently modeled the nominal species distribution throughout the Americas using a low presence threshold (0.08) and found a much larger land proportion occupied by the species than the one reported in this study. Nevertheless, the spatial patterns were consistent, and the difference in km² likely just reflects the reduction in suitable area estimations that can be expected when a much more restrictive threshold (0.7 in this study) is used. This point will be revisited below.

Regarding the importance of the variables used as predictors, our results are in partial agreement with those by Gómez-Llano *et al*. (2025), who highlighted Mean Diurnal Range (BIO2), Mean Temperature of the Wettest Quarter (BIO8), and Precipitation of the Wettest Month (BIO13) as key factors. Yet, our results emphasized Annual Mean Temperature (BIO1) and Mean Temperature of the Warmest Quarter (BIO10), whereas Gómez-Llano *et al*. (2025) underlined other seasonal variables absent from our dataset. These differences reflect methodological choices in variable selection and provide complementary perspectives on *A. fraterculus* distribution. At the morphotype level, most of them were mainly influenced by temperature, with Annual Mean being most relevant for Brazilian-1 and Warmest Quarter for the Peruvian and Ecuadorian cases. Exceptions include Brazilian-3, more affected by Mean Diurnal Range, and the Mexican morphotypes, where precipitation variables were predominant. Perhaps Brazilian-3 is so well adapted to coastal areas with stable temperatures that it is particularly sensitive to wider thermal ranges, while the Mexican case likely reflects stronger seasonal variability in rainfall. Morphotypes at higher elevations, in turn, seem more constrained by warmest-quarter conditions, when temperature extremes are more pronounced.

Ultimately, despite differences in methodology and variables used, all studies show a niche diversification strategy for *A. fraterculus*. When SDM information is broken down into different taxonomic entities (morphotypes), this niche diversification mechanism becomes even more evident. Our results revealed marked contrasts in the distributional extent and ecological preferences of the morphotypes (see Supplementary Figures and Supplementary Table 2). In line with Selivon *et al*. (2022), Brazilian-1 emerged as the most widespread, occupying diverse habitats across several biomes and tolerating a broad range of climatic conditions, which suggests a greater ecological plasticity. Its potential distribution includes southeastern Brazil, Argentine forests, northwestern South America, southern Central America, and eastern Mexico, a significant wider area that the morphotype is currently occupying. In contrast, Brazilian-2 and Brazilian-3 were confined to narrower ranges along the Brazilian coast, where temperature-related variables were the main drivers, highlighting their sensitivity to climatic gradients. Brazilian-2 is narrowly confined to the coastal region around Rio de Janeiro and São Paulo, while Brazilian-3 shows a fragmented distribution along the Brazilian east coast with some inland presence. The Andean, Ecuadorian, Peruvian and Venezuelan morphotypes showed a clear affinity for high-elevation environments in the Northern and Central Andes, with temperature playing a central role in shaping their distributions. Although sharing this elevational niche, each morphotype exhibited subtle differences in range breadth and dominant environmental predictors, reflecting local adaptations. Andean populations occur in Venezuela and northwestern Colombia, whereas the Ecuadorian morphotype is already limited to Ecuador and the northern Andes. The Peruvian morphotype is present in Peru, with potential toward Bolivia and northern Chile, and the Venezuelan occurs across Venezuela, Colombia, and Peru. Finally, the Mexican morphotype is centered in Mexico, Guatemala, and Panama, but models indicate broad suitability across Central America, the United States, Brazil, and the Chaco region. Altogether, these differences underline the importance of considering morphotype-specific ecological requirements and environmental drivers.

In addition to present SDM, in the context of climatic change, it is useful to investigate rates of adaptation and range shifts in pest populations and to predict how populations will react to environmental changes (Porretta *et al*., 2007; Aluja *et al*., 2012; Wei *et al*., 2015) and their ability to invade might be altered (Larson *et al*., 2019; Ma *et al*., 2020). Our projections for the *A. fraterculus* complex indicate that future scenarios are characterized more by localized contractions of high suitability areas (≥0.7) than by broad expansions, with differences between moderate and pessimistic models reflecting changes in intensity rather than geography. The changes in suitability were limited to some portions of the potential distribution, and as expected, the changes deepened over time, so the more distant future scenarios present a deeper change than the near-future ones. Brazilian-1, the most widespread morphotype, is projected to lose large portions of its current range in Brazil, though moderate scenarios suggest expansion into Uruguay and temperate regions of Argentina. Brazilian-2 and Brazilian-3, both restricted coastal forms, are predicted to contract further, while the Andean morphotype remains relatively stable with some inland losses. Ecuadorian and Peruvian distributions shrink considerably, the latter reduced to a narrow coastal strip, and the Venezuelan morphotype is expected to persist mainly in eastern Venezuela after losing much of its current range. By contrast, the Mexican morphotype shows the broadest potential expansion, with high suitability across Central America, parts of the United States, and Brazil, despite losses in Bolivia, Paraguay, and Cuba.

Studies on *Anastrepha* fruit flies (e.g., *A. suspensa*, *A. obliqua*, *A. grandis*, *A. striata*) using CLIMEX and MaxEnt have shown that climate change is likely to expand their potential distribution, particularly in tropical and subtropical regions, with temperature as a main driver and seasonal shifts in suitability (Santos *et al*., 2019; Amat *et al*., 2022; Machado Teixeira *et al*., 2022; Batista Degracia *et al*., 2023; da Silva Santana *et al*., 2023). These models highlight that these fruit flies respond strongly to warming, with range expansions towards higher latitudes and increased environmental suitability in previously marginal areas. The range of thermal tolerance is one of the main factors influencing the geographic distribution of species (Rolandi *et al*., 2018), and therefore differences with congeneric species may be attributable to different thermal thresholds and ecological requirements. At the same time, it has been proposed by Santos (2008) –who also reported a decrease in suitability areas for future scenarios-that water stress under warmer conditions can increase mortality in *A. fraterculus*. Moreover, unlike these other *Anastrepha* species which generally have smaller current ranges, *A. fraterculus* (especially Brazilian-1) already occupies a much broader area, leaving less new territory for expansion but emphasizing the importance of targeted monitoring and management in areas that maintain high suitability. In this line, it is worth noting that Gómez-Llano *et al*. (2025) also informed increases in climatically suitable areas for *A. fraterculus* across all evaluated scenarios using a low presence threshold (0.08), whereas our models — based on a much more restrictive threshold (≥0.7, TSS-optimal)— predict reductions in high-suitability areas for the *AF* complex and most of its morphotypes. This more conservative approach means smaller potential area estimates and that even small decreases in suitability result in reclassification as absence, producing more focused estimates restricted to high-probability occurrence zones. Despite these methodological differences, there is no contradiction: both studies indicate that the future distribution of *A. fraterculus* is likely to shift toward temperate regions across the Americas.

An additional implication is that a reduction in the extent of high-suitability zones should not be interpreted as a decrease in the relevance of the pest. On the contrary, such contractions may concentrate populations in fewer but still highly suitable environments, which can increase local impacts and complicate control strategies. Results of future projections and alpha diversity maps pointed out that the connectivity among *AF* populations in the major fruit-producing regions of South America would be maintained in all scenarios, assuming that these areas will continue their current productivity levels. In this sense, our projections also indicated that the Brazilian-1 morphotype could expand slightly into Argentina. As already reported by Gómez-Llano (2025), the Mexican morphotype is expected to enlarge its high-suitability range in the USA across all scenarios, pointing to additional challenges for future surveillance and management. This highlights the importance of maintaining active monitoring and timely management actions in the areas where favorable conditions persist, as well as anticipating possible shifts in distribution under changing environmental scenarios.

At this point, despite using inferences from multiple algorithms, it is necessary to consider the limitations of SDM and, therefore, those of this study and the limits of its reports. There are several factors that could account for the difference between the potential distribution of a species and the real one, for example, some species could be following the availability of its preferred host (Vera *et al*., 2002), which could also be altered in response to climate change. Other limitations not considered could be biotic interactions (intra– and interspecific) and anthropogenic factors, such as agriculture, urbanization, and land use (Altamirano, 2017). There are well known examples of a species reaching a potentially suitable area but not establishing due to competition with a resident species, like *C. capitata* in Queensland (Vera *et al*. 2002) or some areas of Bolivia (Abad *et al*. 2018), Specifically, regarding resources, given that *A. fraterculus* is highly polyphagous and exhibits metapopulation dynamics with successive processes of colonization-extinction (Freilij *et al*., 2022) and seasonal migration to different hosts, this mitigates the lack of resources, meaning that as long as the different crops are maintained, there will be populations that survive and evolve. Biotic interactions, including competition with other species and potential hybridization among morphotypes, as well as anthropogenic factors, are likely to be more relevant at the local level. Targeted studies addressing these aspects could greatly enhance monitoring efforts and the development of IPM strategies. Natural enemies also play an important role. Gómez-Llano *et al*. (2025) evaluated the joint distribution of *A. fraterculus* and two parasitoids (*Doryctobracon areolatus* and *Diachasmimorpha longicaudata*), reporting expansions in suitable areas for their establishment under future scenarios. Their approach underscores the significance of considering both pest and natural enemy distributions in the planning of effective IPM strategies. Studies examining the evolution of thermal thresholds for all morphotypes will contribute to understanding their experimental response to rising temperatures resulting from climate change. Similarly, genomic studies that unravel the genetic basis of local adaptation of morphotypes, as well as genomic offset, will be key to projecting the response of the species and regional and local risks. Addressing these ecological, physiological, and genomic aspects in future studies will be crucial for improving predictive models and informing more effective and context-specific IPM strategies for *A. fraterculus* across the Americas.

## Conclusions

The focus of this study was the distribution of the South American fruit fly, *Anastrepha fraterculus*, a species that comprises at least eight morphotypes. We provide a comprehensive assessment of the current and future geographic distribution of the nominal species and all its morphotypes under different scenarios of climate change. The modeling was robust, with trustworthy projections showing some areas that will remain highly suitable for some of the morphotypes and others that will not, which would lead to a significant migration of the flies toward temperate regions. This will generate new interactions between the different morphotypes, their plant hosts and other species that may constrain the establishment of *A. fraterculus*, as competitors, predators or parasites. This study provides a valuable tool in the context of monitoring and implementing IMP strategies, serving as a reliable reference and opening new horizons for research, both in wide and local geographical areas.

## Acknowledgments

This work was supported by fundings from the Agencia Nacional de Promoción Científica y Técnica (ANPCYT; PICT 2018-02567 granted to Dr M. I. Remis) and Universidad de Buenos Aires (UBA; UBACYT 20020170100270BA to J.C.V.). D.F. holds a doctoral fellowship from the Consejo Nacional de Investigaciones Científicas y Técnicas (CONICET, Argentina). J.C.V is a researcher at CONICET.

## Disclosure

The authors declare no competing interests.

## Supplementary Tables captions

**Supplementary Table 1.** Geographical coordinates morphotype, and original source citation for each *Anastrepha fraterculus* recording.

**Supplementary Table 2.** Areas with high climatic suitability (calculated from the binary 0.7 TSS threshold projection) for *Anastrepha fraterculus* under current and future climate scenarios. f1= MRI-ESM2-SSP3-7.0 for 2021-2040; f2= MRI-ESM2-SSP3-7.0 for 2041-2060; f3= UKESM1-0-LL-SSP5-8.5 for 2021-2040; UKESM1-0-LL-SSP5-8.5 for 2041-2060.

Data are sorted for the whole *AF* complex and for each morphotype, in separate sheets for country and for ecoregion.

Sheet 2: *A. fraterculus* complex by country.

Sheet 3: *A. fraterculus* complex by ecoregion.

Sheet 4: Andean morphotype by country.

Sheet 5: Andean morphotype by ecoregion.

Sheet 6: Brazilian-1 morphotype by country.

Sheet 7: Brazilian-1 morphotype by ecoregion.

Sheet 8: Brazilian-2 morphotype by country.

Sheet 9: Brazilian-2 morphotype by ecoregion.

Sheet 10: Brazilian-3 morphotype by country.

Sheet 11: Brazilian-3 morphotype by ecoregion.

Sheet 12: Ecuadorian morphotype by country.

Sheet 13: Ecuadorian morphotype by ecoregion.

Sheet 14: Mexican morphotype by country.

Sheet 15: Mexican morphotype by ecoregion.

Sheet 16: Peruvian morphotype by country.

Sheet 17: Peruvian morphotype by ecoregion.

Sheet 18: Venezuelan morphotype by country.

Sheet 19: Venezuelan morphotype by ecoregion.

**Supplementary Figure 1:**
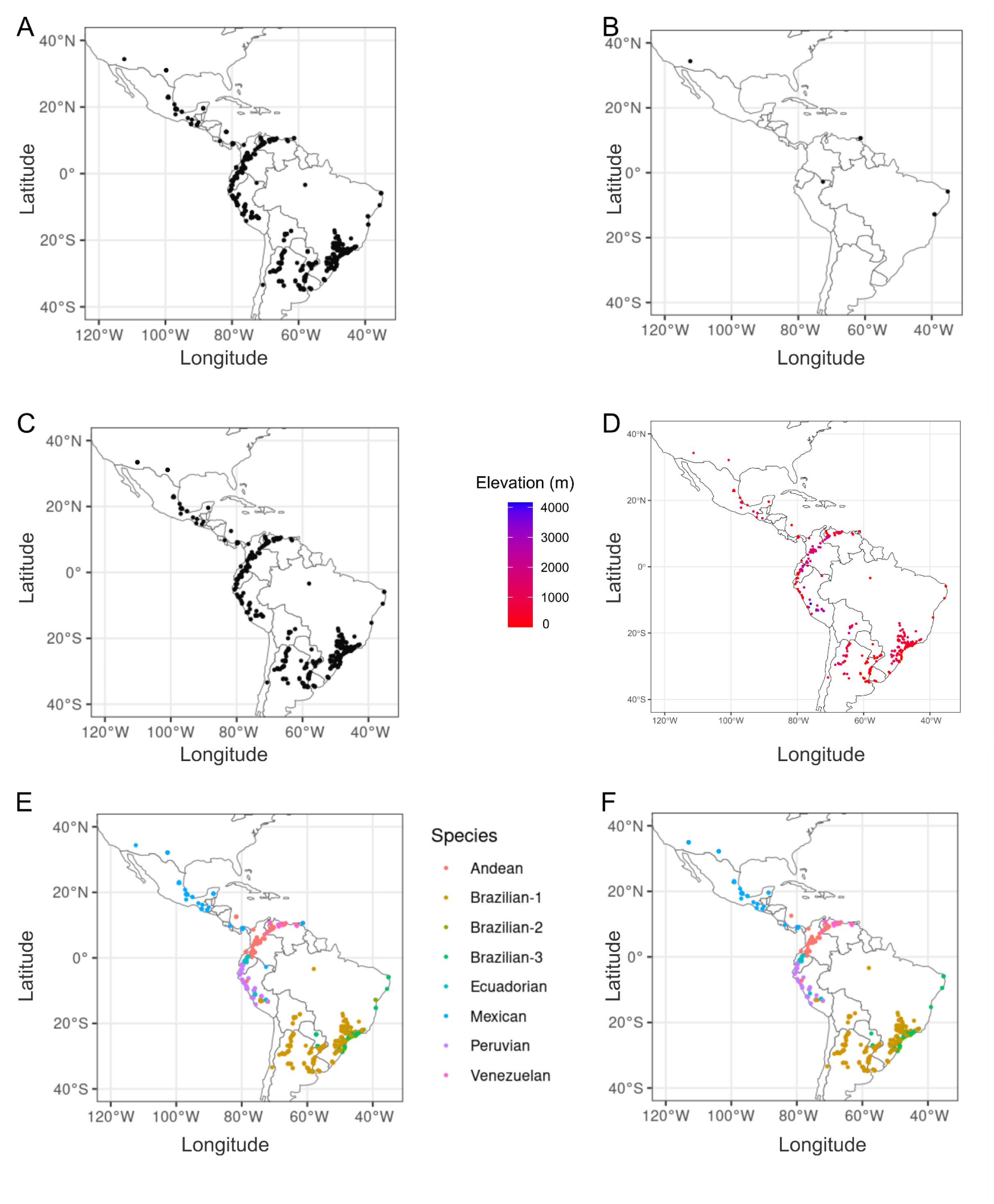
Occurrence points cleaning. **A.** 1412 initial points of occurrence for the *Anastrepha fratercu/us* complex throughout the Americas. **B.** Putative outliers detected using the CoordinateCleaner package. **C.** Post-filtering occurrence points. **D.** Elevation of post-filtered occurrence points. **E.** Occurrence points with pre-filtered morphotype information. **F.** Occurrence points for each morphotype post filtering.

**Supplementary Figure 2:**
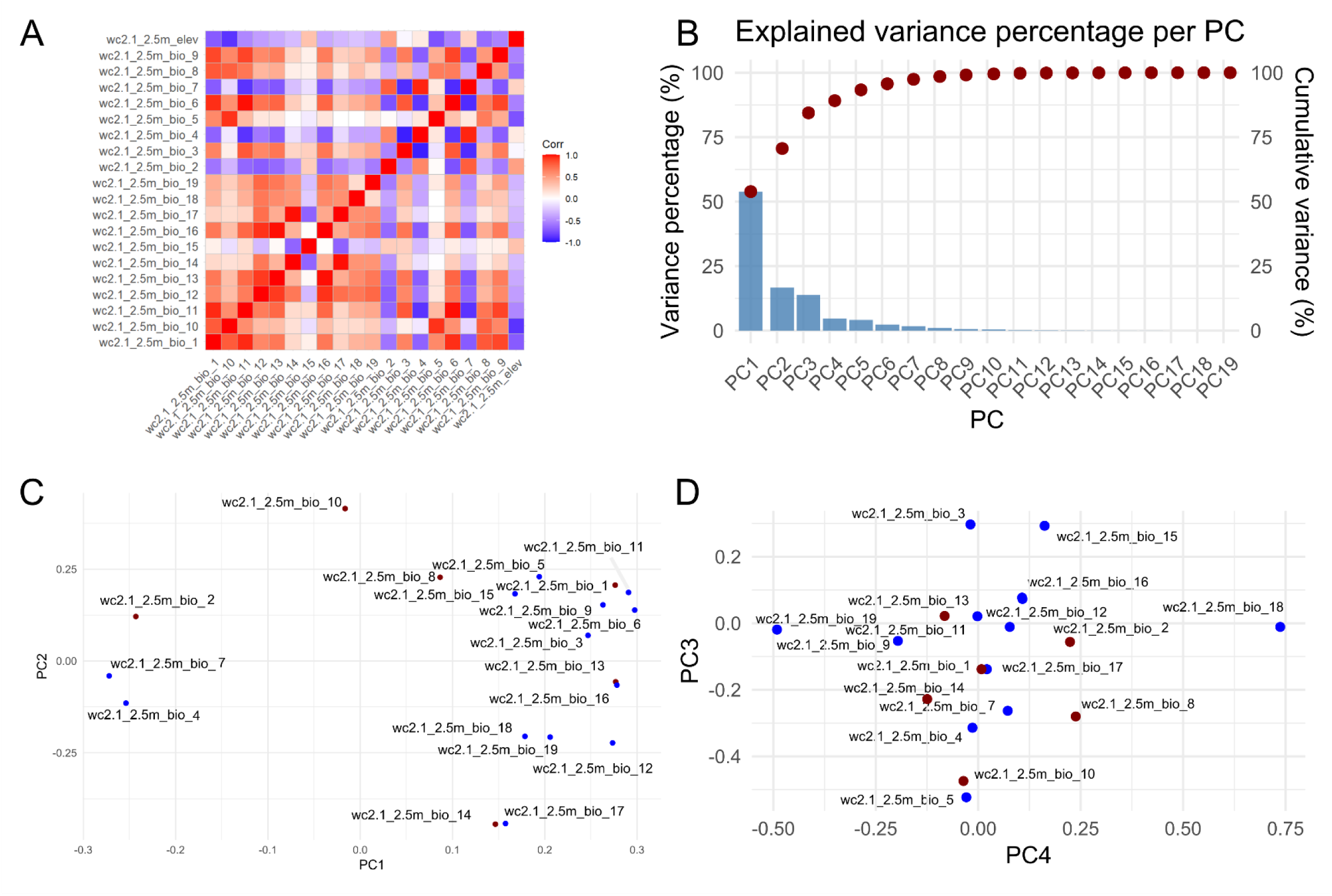
Environmental data filtering. **A.** Correlation heatmap for WorldClim 2.5 arc min bioclimatic variables and elevation. **B.** Explained variance percentage for the Principal Component Analysis (PCA) of bioclimatic variables. **C.** PCA variable contributions to PC 1-2. **D.** PCA variable contributions to PC 3-4. C-D dark red points represent the selected variables for SOM; blue dots filtered variables.

**Supplementary Figure 3:**
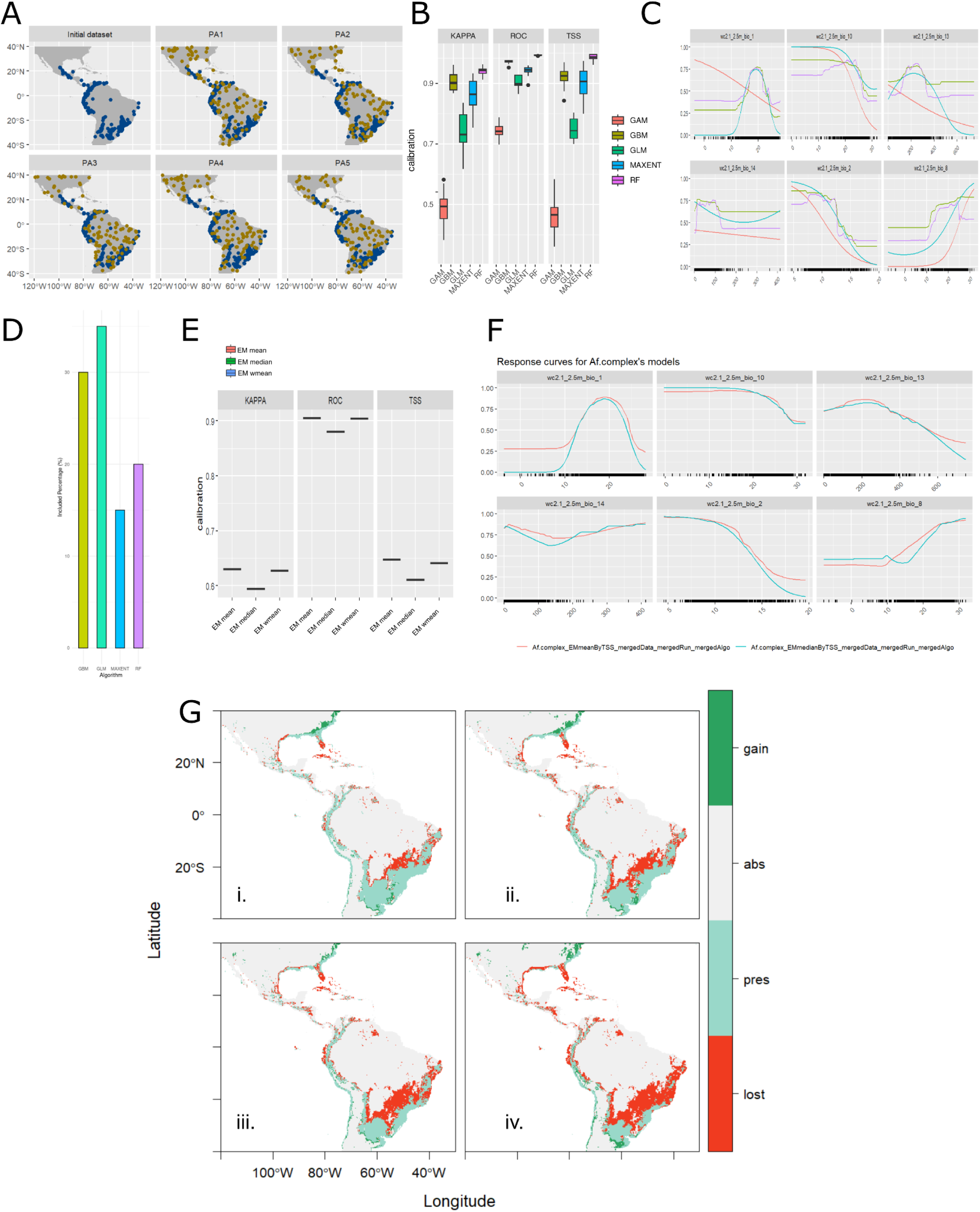
Modelling details for the *Anastrepha fratercu/us* complex. **A.** Pseudo-absences points for each repetition. Blue points=presences; yellow points= pseudo-absences. **B.** Calibration results for cross validation per algorithm. **C.** Variables response curves for each algorithm. **D.** Contribution of each algorithm to the ensemble model. **E.** Ensemble model evaluation metrics. F. Response curves for the ensemble model. **G.** Range change map under different climate change scenarios: i= MRI-**ESM2-SSP3-7.0** for **2021-2040;** ii= **MRI-ESM2-SSP3-7.0 for 2041-2060;** iii= **UKESM1-0-LL-SSP5-8.5** for **2021-2040; iv= UKESM1-0-LL-SSP5-8.5 for 2041-2060.**

**Supplementary Figure 4:**
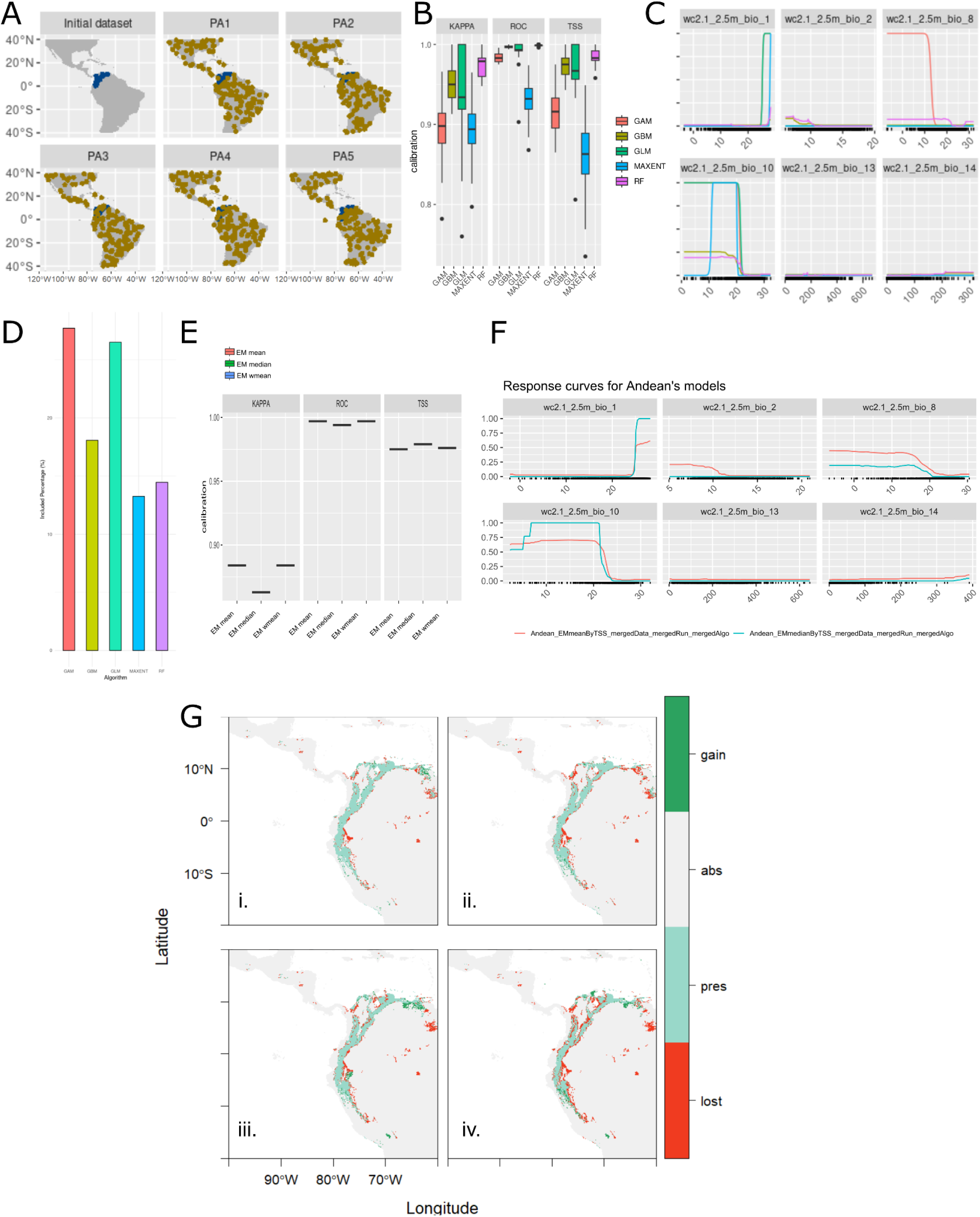
Modelling details for the Andean morphotype. **A.** Pseudo-absences points for each repetition. Blue points=presences; yellow points= pseudo-absences. **B.** Calibration results for cross validation per algorithm. **C.** Variables response curves for each algorithm. **D.** Contribution of each algorithm to the ensemble model. **E.** Ensemble model evaluation metrics. **F.** Response curves for the ensemble model. **G.** Range change map under different climate change scenarios: **i=** MRI-ESM2-SSP3-7.0 **for 2021-2040;** ii= **MRI-ESM2-SSP3-7.0 for 2041-2060;** iii= **UKESM1-0-LL-SSP5-8.5** for **2021-2040; iv= UKESM1-0-LL-SSP5-8.5 for 2041-2060.**

**Supplementary Figure 5:**
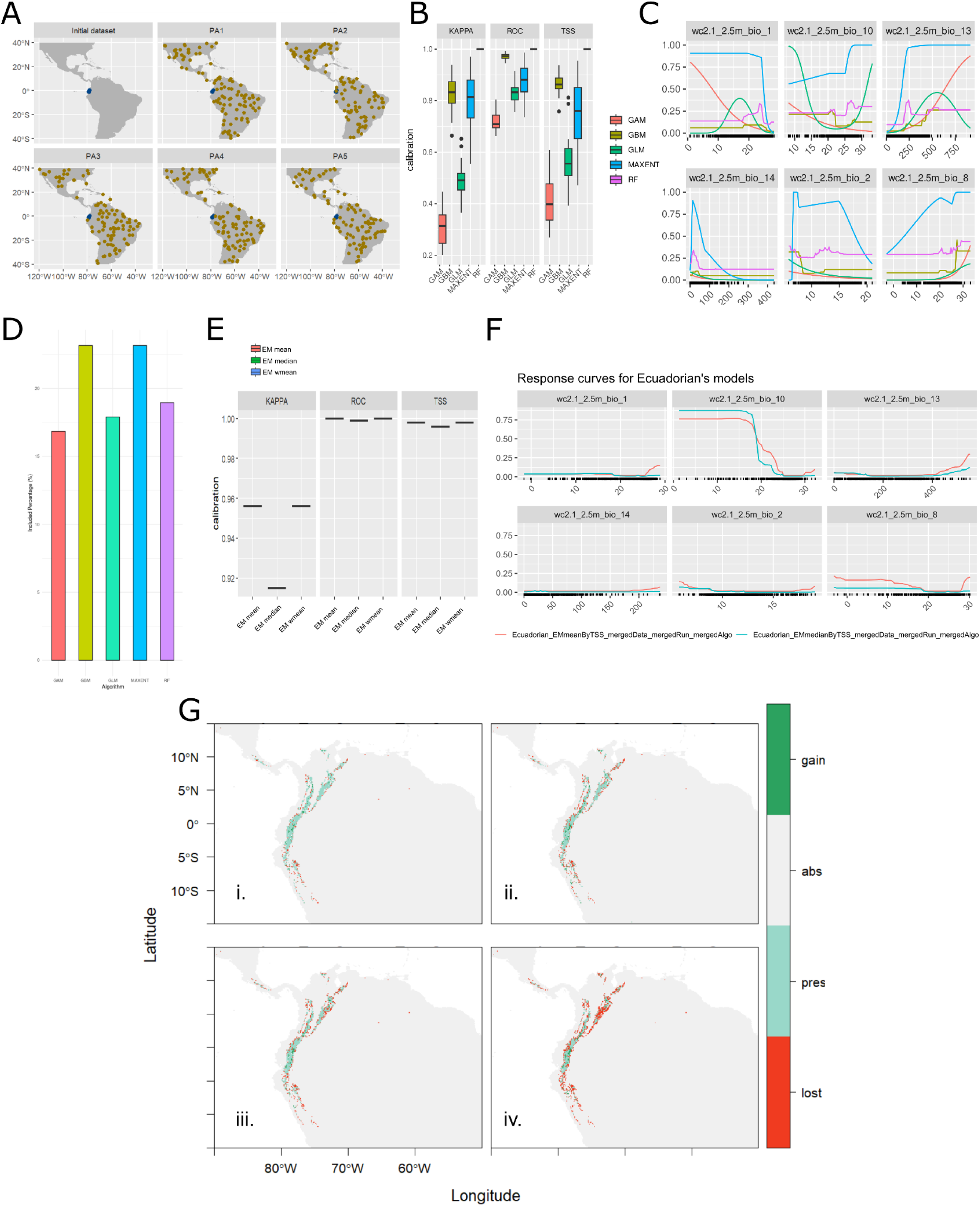
Modelling details for the Ecuadorian morphotype. **A.** Pseudo-absences points for each repetition. Blue points=presences; yellow points= pseudo-absences. **B.** Calibration results for cross validation per algorithm. **C.** Variables response curves for each algorithm. **D.** Contribution of each algorithm to the ensemble model. **E.** Ensemble model evaluation metrics. **F.** Response curves for the ensemble model. **G.** Range change map under different climate change scenarios: i= MRI-ESM2-SSP3-7.0 for 2021-2040; ii= MRI-ESM2-SSP3-7.0 for 2041-2060; iii= UKESM1-0-LL-SSP5-8.5 for 2021-2040; iv= UKESM1-0-LL-SSP5-8.5 for 2041-2060.

**Supplementary Figure 6:**
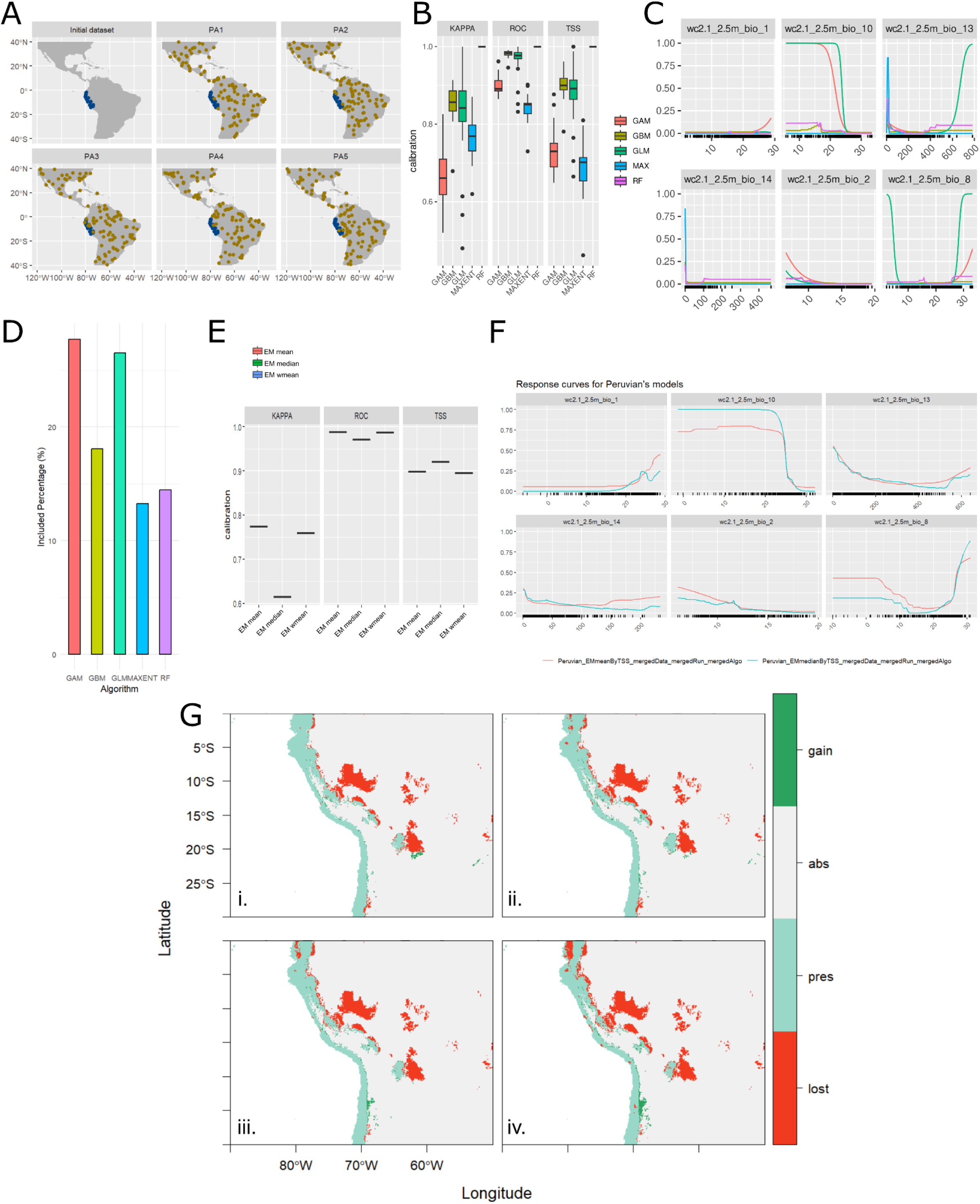
Modelling details for the Peruvian morphotype. **A.** Pseudo-absences points for each repetition. Blue points=presences; yellow points= pseudo-absences. **B.** Calibration results for cross validation per algorithm. **C.** Variables response curves for each algorithm. **D.** Contribution of each algorithm to the ensemble model. **E.** Ensemble model evaluation metrics. **F.** Response curves for the ensemble model. **G.** Range change map under different climate change scenarios: i= MRI-ESM2-SSP3-7.0 for 2021-2040; ii= MRI-ESM2-SSP3-7.0 for 2041-2060; iii= UKESM1-0-LL-SSP5-8.5 for 2021-2040; iv= UKESM1-0-LL-SSP5-8.5 for 2041-2060.

**Supplementary Figure 7:**
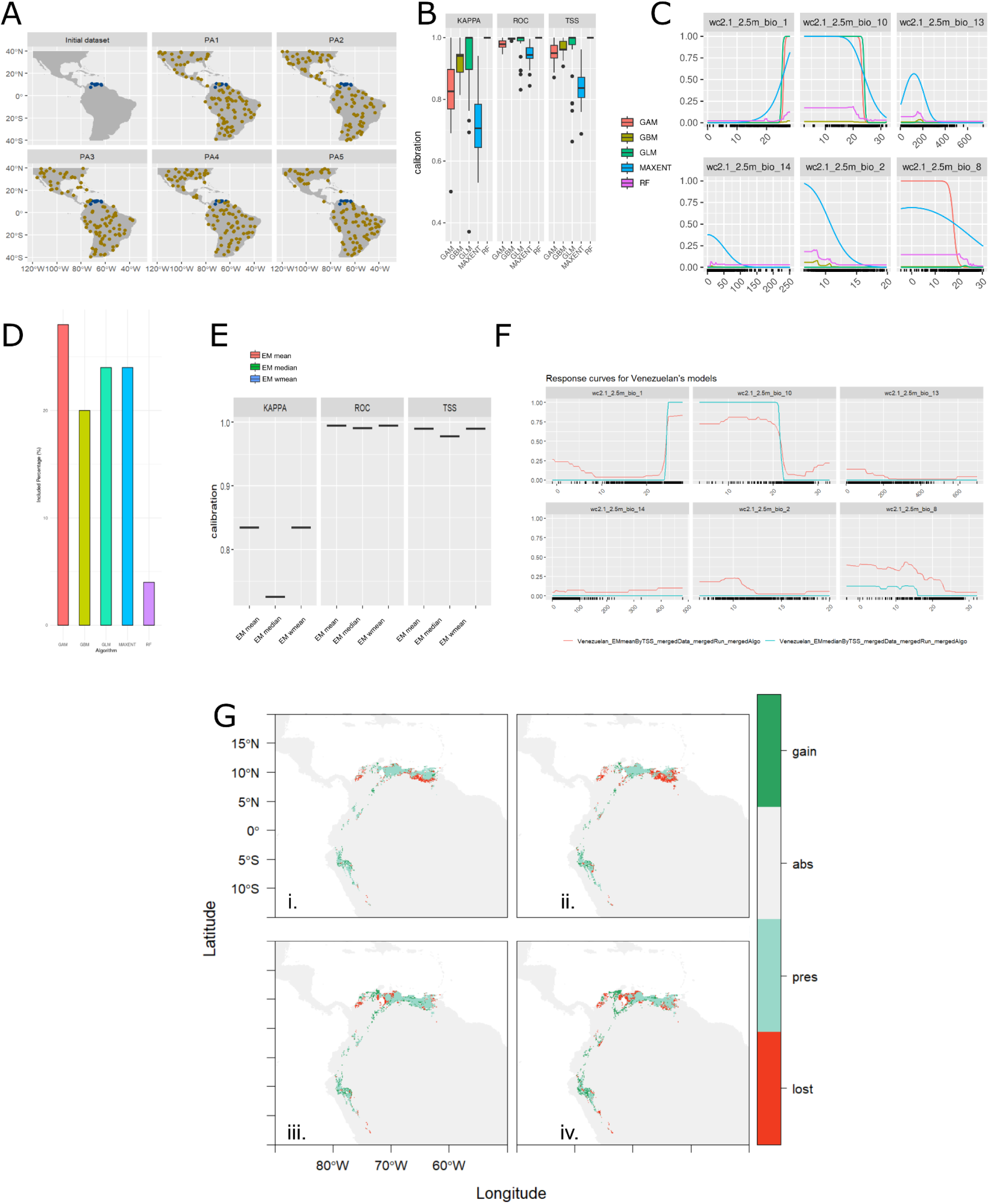
Modelling details for the Venezuelan morphotype. **A.** Pseudo-absences points for each repetition. Blue points-presences; yellow points= pseudo-absences. B. Calibration results for cross validation per algorithm. **C.** Variables response curves for each algorithm. **D.** Contribution of each algorithm to the ensemble model. **E.** Ensemble model evaluation metrics. **F.** Response curves for the ensemble model. G. Range change map under different climate change scenarios: i= MRI-ESM2-SSP3-7.0 for 2021-2040; ii= MRI-ESM2-SSP3-7.0 for 2041-2060; iii= UKESM1-0-LL-SSP5-8.5 for 2021-2040; iv= UKESM1-0-LL-SSP5-8.5 for 2041-2060.

**Supplementary Figure 8:**
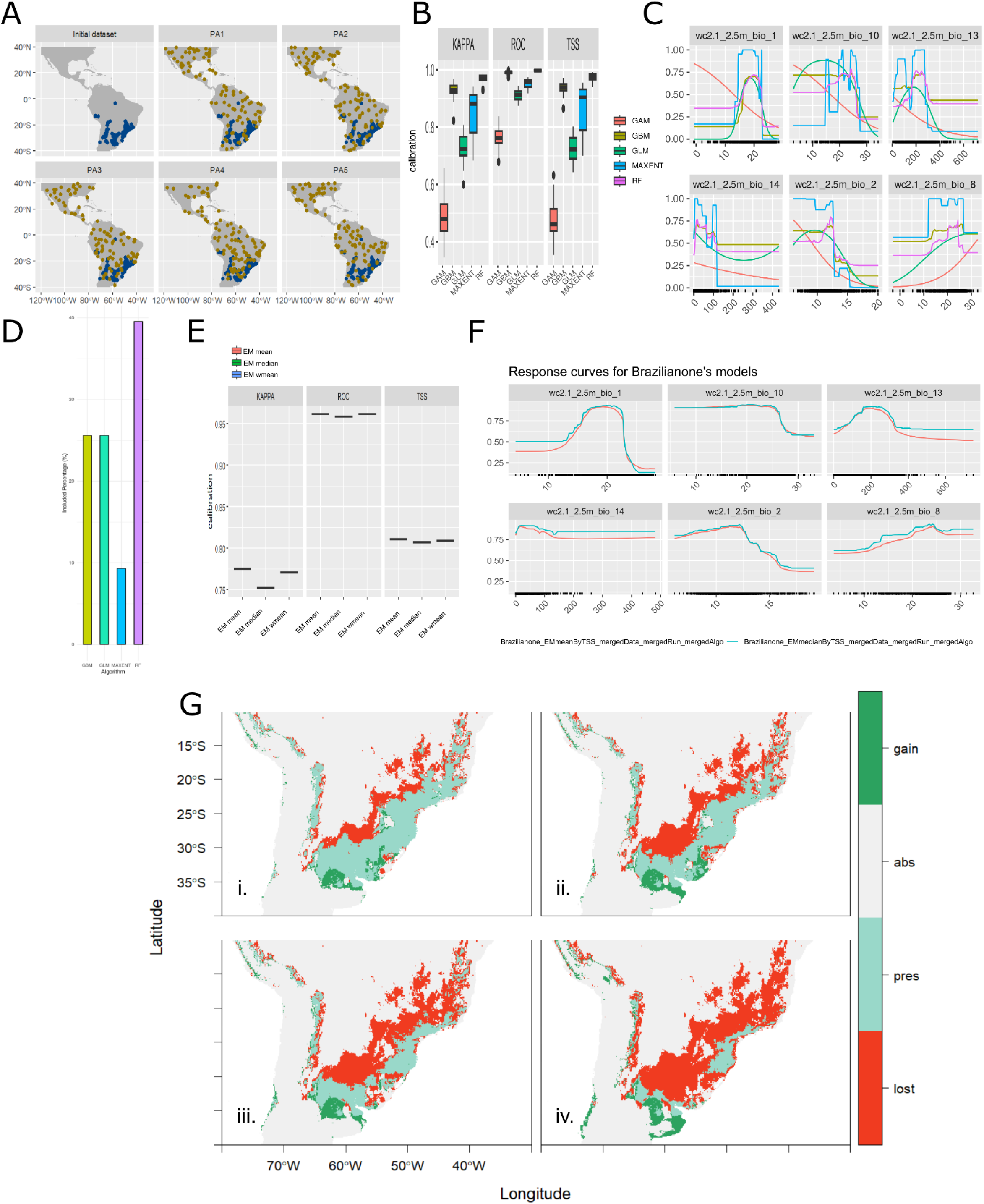
Modelling details for the Brazilian-1 morphotype. **A.** Pseudo-absences points for each repetition. Blue points-presences; yellow points= pseudo-absences. B. Calibration results for cross validation per algorithm. **C.** Variables response curves for each algorithm. **D.** Contribution of each algorithm to the ensemble model. **E.** Ensemble model evaluation metrics. **F.** Response curves for the ensemble model. G. Range change map under different climate change scenarios: i= MRI-ESM2-SSP3-7.0 for 2021-2040; ii= MRI-ESM2-SSP3-7.0 for 2041-2060; iii= UKESM1-0-LL-SSP5-8.5 for 2021-2040; iv= UKESM1-0-LL-SSP5-8.5 for 2041-2060.

**Supplementary Figure 9:**
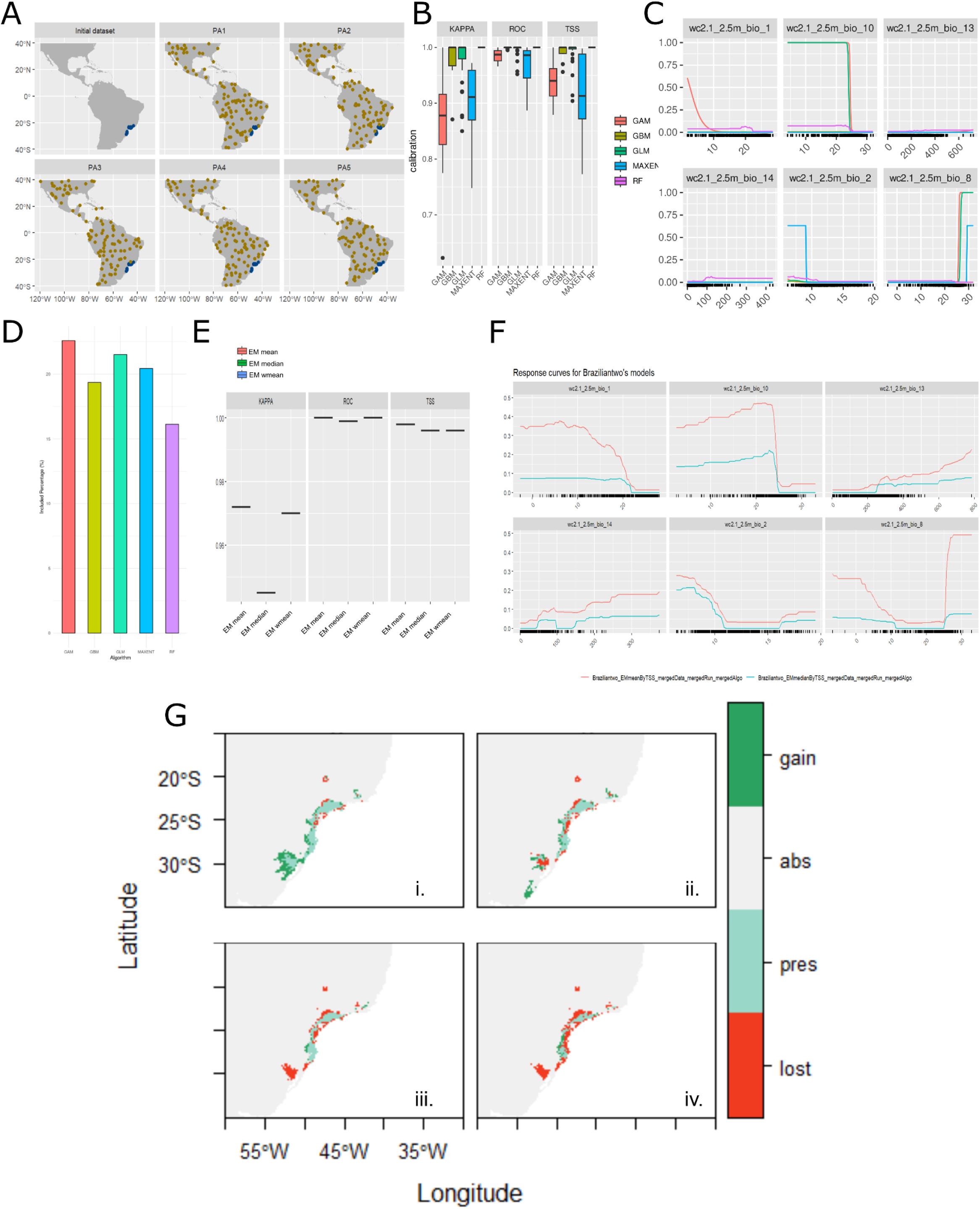
Modelling details for the Brazilian-2 morphotype. **A.** Pseudo-absences points for each repetition. Blue points-presences; yellow points= pseudo-absences. B. Calibration results for cross validation per algorithm. **C.** Variables response curves for each algorithm. **D.** Contribution of each algorithm to the ensemble model. **E.** Ensemble model evaluation metrics. **F.** Response curves for the ensemble model. G. Range change map under different climate change scenarios: i= MRI-ESM2-SSP3-7.0 for 2021-2040; ii= MRI-ESM2-SSP3-7.0 for 2041-2060; iii= UKESM1-0-LL-SSP5-8.5 for 2021-2040; iv= UKESM1-0-LL-SSP5-8.5 for 2041-2060.

**Supplementary Figure 10:**
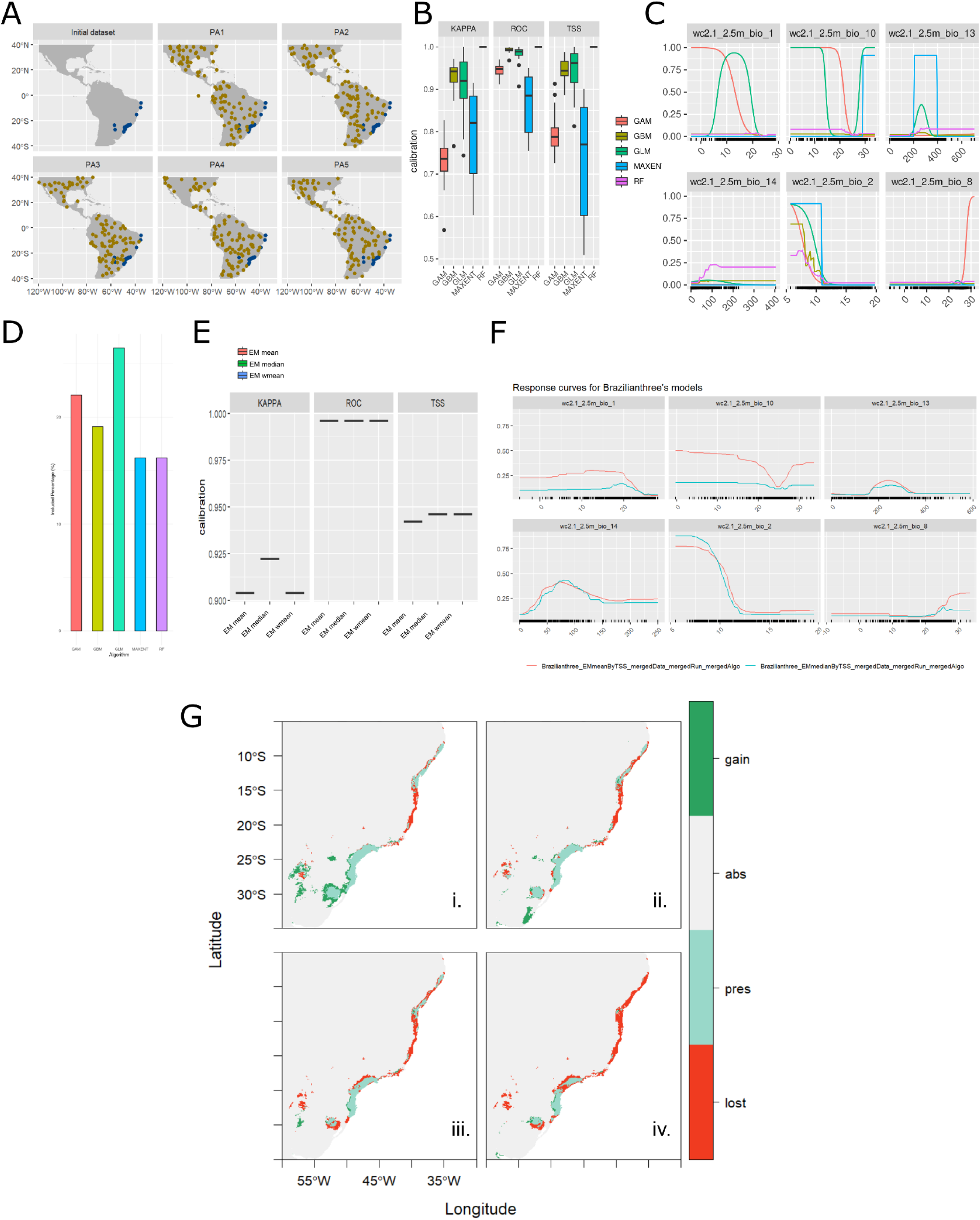
Modelling details for the Brazilian-3 morphotype. **A.** Pseudo-absences points for each repetition. Blue points-presences; yellow points= pseudo-absences. B. Calibration results for cross validation per algorithm. **C.** Variables response curves for each algorithm. **D.** Contribution of each algorithm to the ensemble model. **E.** Ensemble model evaluation metrics. **F.** Response curves for the ensemble model. G. Range change map under different climate change scenarios: i= MRI-ESM2-SSP3-7.0 for 2021-2040; ii= MRI-ESM2-SSP3-7.0 for 2041-2060; iii= UKESM1-0-LL-SSP5-8.5 for 2021-2040; iv= UKESM1-0-LL-SSP5-8.5 for 2041-2060.

**Supplementary Figure 11:**
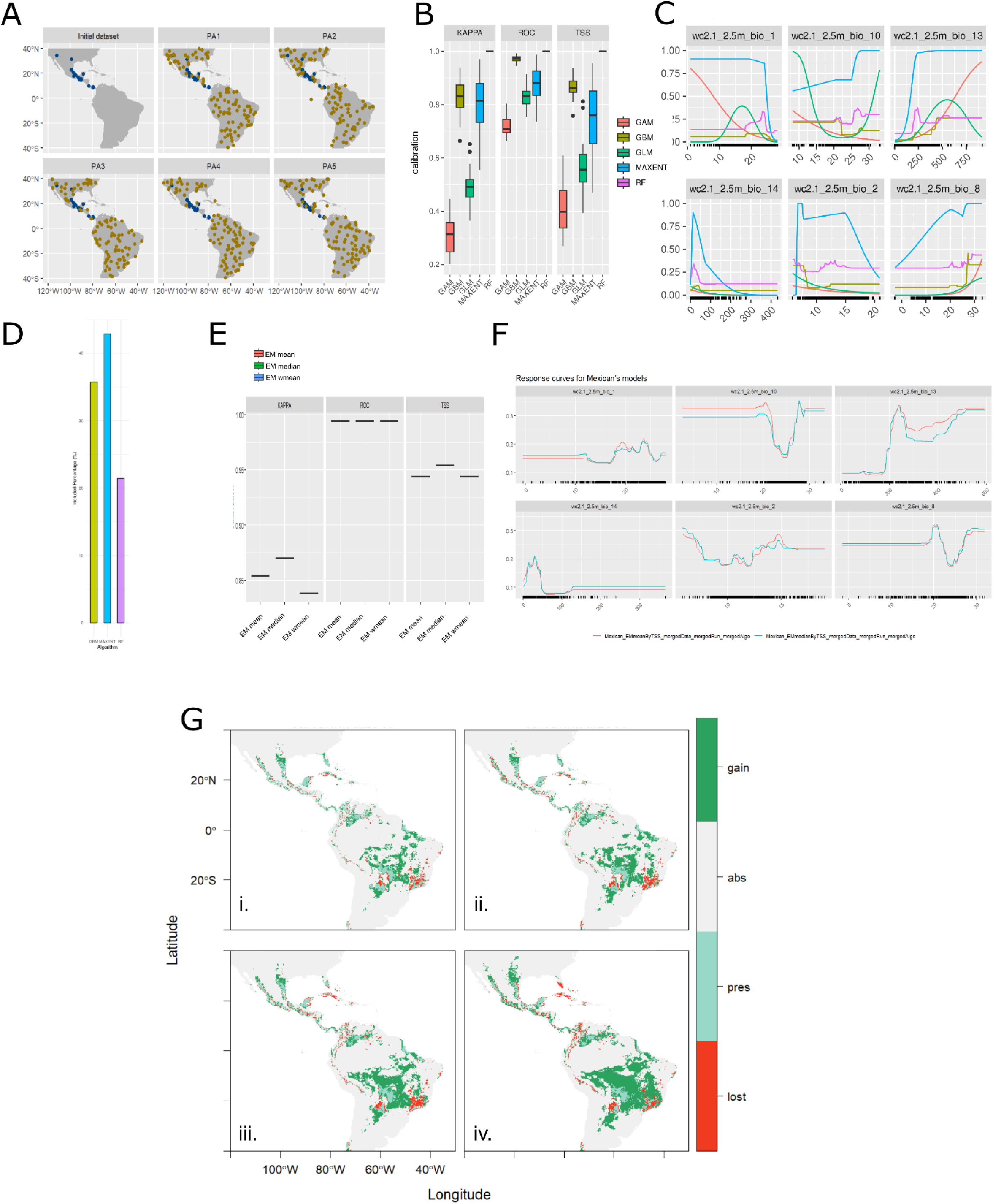
Modelling details for the Mexican morphotype. **A.** Pseudo-absences points for each repetition. Blue points=presences; yellow points-pseudo-absences. **B.** Calibration results for cross validation per algorithm. **C.** Variables response curves for each algorithm. **D.** Contribution of each algorithm to the ensemble model. **E.** Ensemble model evaluation metrics. **F.** Response curves for the ensemble model. G. Range change map under different climate change scenarios: i= MRI-ESM2-SSP3-7.0 for 2021-2040; ii= MRI-ESM2-SSP3-7.0 for 2041-2060; iii= UKESM1-0-LL-SSP5-8.5 for 2021-2040; iv= UKESM1-0-LL-SSP5-8.5 for 2041-2060.

**Supplementary Figure 12:**
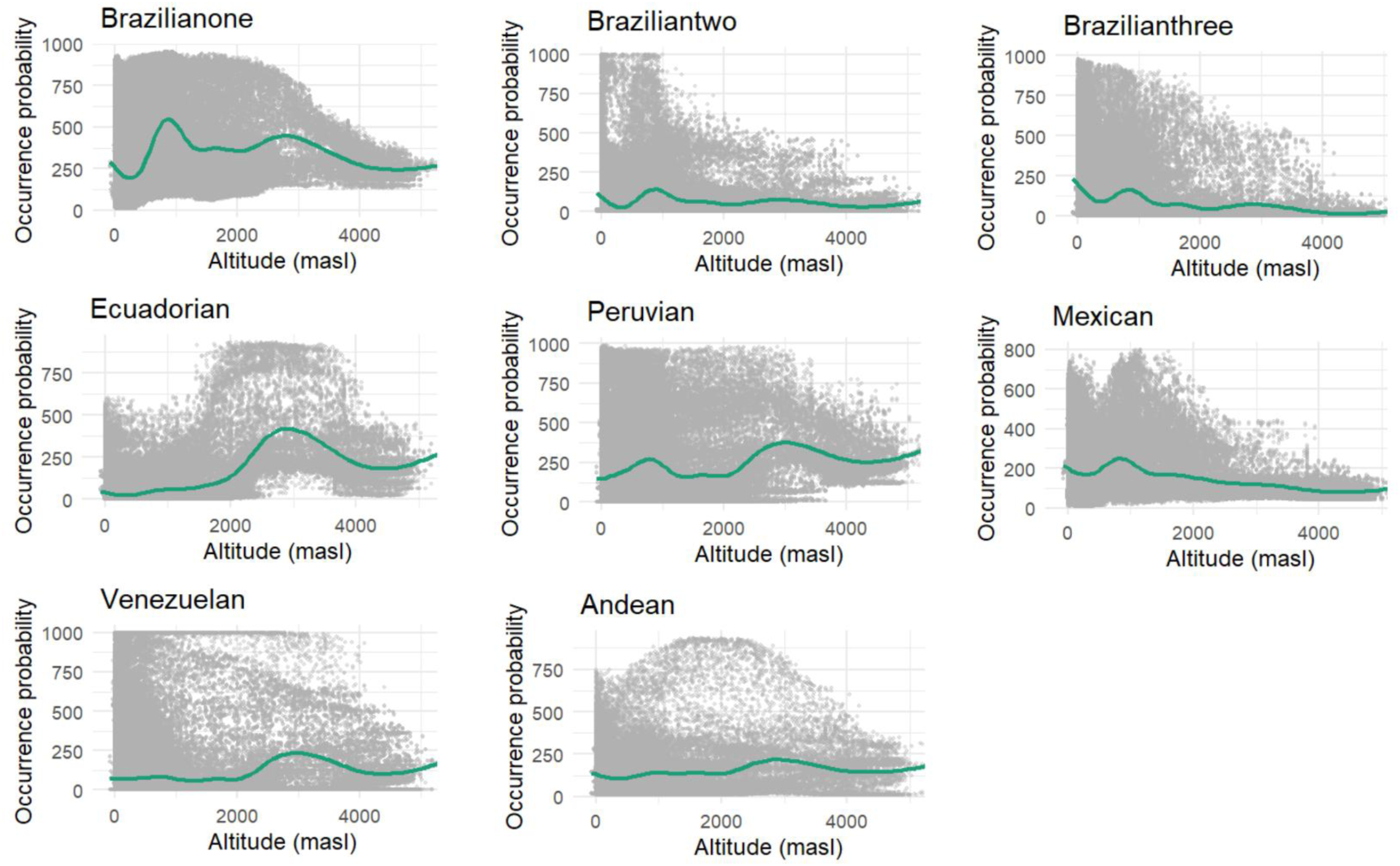
Ocurrence probability with elevation per morphotype of the *AF* complex.

## Notes

### Competing Interest Statement

The authors have declared no competing interest.

